# ClairS: a deep-learning method for long-read somatic small variant calling

**DOI:** 10.1101/2023.08.17.553778

**Authors:** Zhenxian Zheng, Junhao Su, Lei Chen, Yan-Lam Lee, Tak-Wah Lam, Ruibang Luo

## Abstract

Identifying somatic variants in tumor samples is a crucial task, which is often performed using statistical methods and heuristic filters applied to short-read data. However, with the increasing demand for long-read somatic variant calling, existing methods have fallen short. To address this gap, we present ClairS, the first deep-learning-based, long-read somatic small variant caller. ClairS was trained on massive synthetic somatic variants with diverse coverages and variant allele frequencies (VAF), enabling it to accurately detect a wide range of somatic variants from paired tumor and normal samples. We evaluated ClairS using the latest Nanopore Q20+ HCC1395-HCC1395BL dataset. With 50-fold/25-fold tumor/normal, ClairS achieved a 93.01%/86.86% precision/recall rate for Single Nucleotide Variation (SNVs), and 66.54%/66.89% for somatic insertions and deletions (Indels). Applying ClairS to short-read datasets from multiple sources showed comparable or better performance than Strelka2 and Mutect2. Our findings suggest that improved read phasing enabled by long-read sequencing is key to accurate long-read SNV calling, especially for variants with low VAF. Through experiments across various coverage, purity, and contamination settings, we demonstrated that ClairS is a reliable somatic variant caller. ClairS is open-source at https://github.com/HKU-BAL/ClairS.

## Introduction

Analysis of cancer genomes that identify and characterize somatic variants has enabled a better understanding of tumor progression^1^ and led to precision oncology^2^. Identifying somatic variants, however, remains challenging due to intra– and inter-tumor heterogeneity, which often leads to low VAF, and confounding factors, including sequencing artifacts, inadequate sequencing coverage, and normal contamination^3^. Endeavors were made to address these challenges and maximize sensitivity and accuracy in identifying somatic variants using next-generation sequencing (NGS) short-reads^4–13^. However, constrained by read length, short reads have limited variant discovery capability in hard-to-map genomic regions, such as homopolymers and segmental duplications. This problem is expected to be alleviated through long-read sequencing^14^. Oxford Nanopore Technologies (ONT) is among the leading long-read sequencing technologies and offers miniaturized sequencing devices and fast sample-to-data turnaround, which is a credible step towards democratizing sequencing by drastically reducing the cost of carrying out sequencing experiments. ONT raw reads were reported to have an error rate of 3-15% in the past^15^. This was reduced to 1% or lower using ONT’s latest Q20+ chemistry^16^. The gap is still significant, however, compared with NGS short-reads, which have an average error rate at 0.1%^17^, making the somatic variant callers once designed for short reads practically unworkable for ONT long-reads.

Germline variants are often considered to be easier to correctly identify than somatic variant calling. The first attempt to call germline small variants using noisy ONT long-reads was made by Clairvoyante in 2018^18^. The work was enabled using 1) a deep neural network, which was first used for variant calling by DeepVariant^19^; and 2) the high-quality known truth variants in GIAB reference samples for neural network model training^20^. Subsequent works by Clairvoyante including Clair^21^ and Clair3^22^, introduced optimized network input, network output, network architecture, and workflow designs to make the best out of noisy ONT data for germline small variant calling. Both Clair3 and a pipeline named PEPPER-Margin-DeepVariant^23^ (DeepVariant), which is also designed for ONT long-read germline small-variant calling, have demonstrated better single nucleotide polymorphism (SNP)-calling performance than using the same coverage of Illumina short reads. However, while solutions are ready for ONT long-read germline small-variant calling, there has been no caller available for ONT long-read somatic small-variant calling. We note that ONT long-read somatic SV (structural variant) callers, including Sniffles2^24^ and Nanomonsv^25^, which were developed in the past year called for the development of a small variant caller to complete the ONT long-read somatic variant-calling workflow.

Unfortunately, some designs critical to ONT long-read germline variant calling are not applicable to somatic variant calling. First, in their network output, both Clair3 and DeepVariant apply a strong diploid genome assumption. Clair3 uses a 21-genotype output, which is a two-combination of A, C, G, T, insertion, and deletion^22^. DeepVariant uses a three-category output that includes hom-ref (homozygous reference), het (heterozygous), and hom-alt (homozygous alternative)^23^. Both Clair3 and DeepVariant are classification models that use observed allele frequency of alternative alleles as network input, and output the category that represents the expected allele frequency of a variant (e.g., allele frequency 0, 0.5, 1, and 0.5/0.5 for genotype 0/0, 0/1, 1/1, and 1/2, respectively). However, somatic variants have VAF ranging continuously from 0 to 1. Without a certain ploidy, somatic variant candidates have no expected allele frequency for a model to test against. Thus, a new design is required. As an example, a new design could use a regression model to derive VAF directly, or a classification model to simply determine whether a candidate is a somatic variant or not and infer VAF subsequently. Second, the seven standard GIAB reference samples HG001–HG007 provide approximately 25 million truth germline variants^20^, which are critical for the model training of any state-of-the-art, deep-learning-based germline variant callers. However, in terms of known truth somatic variants, only the HCC1395–HCC1395BL (a human triple-negative breast cancer cell line and a normal cell line derived from the B lymphocytes of the same donor, hereafter referred to as HCC1395/BL) tumor-normal pair was published by the Somatic Mutation Working Group of the SEQC2 (Sequencing Quality Control Phase II) consortium^3^. It contains only 39,560 SNVs and 1,922 Indels, which is orders of magnitude fewer than the available truth germline variants, and far from enough for deep neural-network model training. In the absence of adequate real tumor-normal samples, one could think of synthetic data as a solution. Bamsurgeon spiked somatic variants into sequencing reads to mimic a tumor^26^. This was a successful method for short reads, but it doesn’t work for long reads for two reasons: 1) Illumina short reads are read one base at a time, whereas ONT long reads are read as the signals of a sliding 5-mer or 6-mer window. That is, a spike-in variant also changes the signals of the adjacent bases. Bases can be base-called from signals but cannot be authentically turned back into signals, so the spike-in method cannot be applied to long reads; and 2) Bamsurgeon does not apply the point that a somatic variant is usually found in only one haplotype (either maternal or paternal) but is missing in another. The use of long reads over short reads for somatic variant calling makes sense only if the long-read advantage of haplotyping (also called phasing) is utilized. All things considered, a new method for synthesizing long-read data to contain abundant plausible somatic variants is needed.

In this study, we present Clair-Somatic (ClairS), the first somatic small variant caller for ONT long reads, which is named after its germline variant caller predecessors. ClairS draws on the successful experience of the Clair series, and uses a new network output and a new workflow design to address the continuous VAF space of somatic variants. By considering two different samples, A and B, as tumor and normal, respectively, and deeming a germline variant specific to A as a somatic variant against B, we devised a data synthetic strategy that uses only the real long reads of GIAB reference samples with known germline variants, but can simulate somatic variants of any tumor purity, sequencing coverage (lower than the data source), and level of normal contamination. The strategy can theoretically produce an infinite number of somatic variants for model training. We show results to highlight how phasing improves somatic variant-calling performance on long reads. To leverage more remote alignment information that is computationally impractical to include in the network input, we devised a post-processing step that searches for ancestral haplotype support for any somatic variant candidates. This step removed a considerable amount of false positive calls in our experiments. For benchmarking, we sequenced in total 75-fold HCC1395 and 45-fold HCC1395BL ONT Q20+ long-reads (data deposited to NCBI SRA), and used the truth somatic variants provided by the SEQC2 consortium^3^. With 50-fold/25-fold tumor/normal, ClairS achieved 86.86%/93.01% recall/precision rate SNVs, and 66.89%/66.54% for somatic Indels when targeting VAF ≥0.05. For variants with VAF ≥0.2, the numbers go up to 94.65%/96.63% for SNVs, and 73.22%/77.35% for somatic Indels. We also show the performance of ClairS at different tumor/normal coverages, tumor purity and normal contamination. ClairS is designed for ONT long reads, but the whole method is also applicable to Illumina short reads. This versatility allowed us to benchmark ClairS against state-of-the-art short-read somatic variant callers considering that there is no other long-read somatic variant caller to benchmark against. The results show that ClairS performed comparably or slightly better than the current heuristic-based and deep-learning-based callers on short reads.

## Results

### The ClairS method

The ClairS method is elaborated in the Method section. Texts in bold in this and the following two paragraphs can be found as subsection titles in the Method section. In the use of deep-learning for long-read somatic variant calling, ClairS has made breakthroughs in **Training data synthesis** and the **ClairS workflow and design**.

Regarding training data synthesis, Figure 1a shows the workflow for **Generating synthetic tumor and synthetic normal**. As we required sampling without replacement, and in view of the common practice that a tumor sample having higher coverage than its matching normal, we gave **Coverage advice for the source data**. Details of how homozygous and heterozygous germline variants in different samples are converted into somatic variants are given in **Deriving multiple categories of variants from a synthetic tumor/normal pair**. Setups to avoid practical concerns and limitations of the data synthesis method are given in **Other details about the variants selected for model training**. Our data synthesis method is based on the observation that an authentic somatic variant is usually found in the reads of a single haplotype (depicted in Extended Data Figure 1a). Also, we found that **Phasing information enhances somatic variant calling performance** in ClairS.

**Figure 1.**
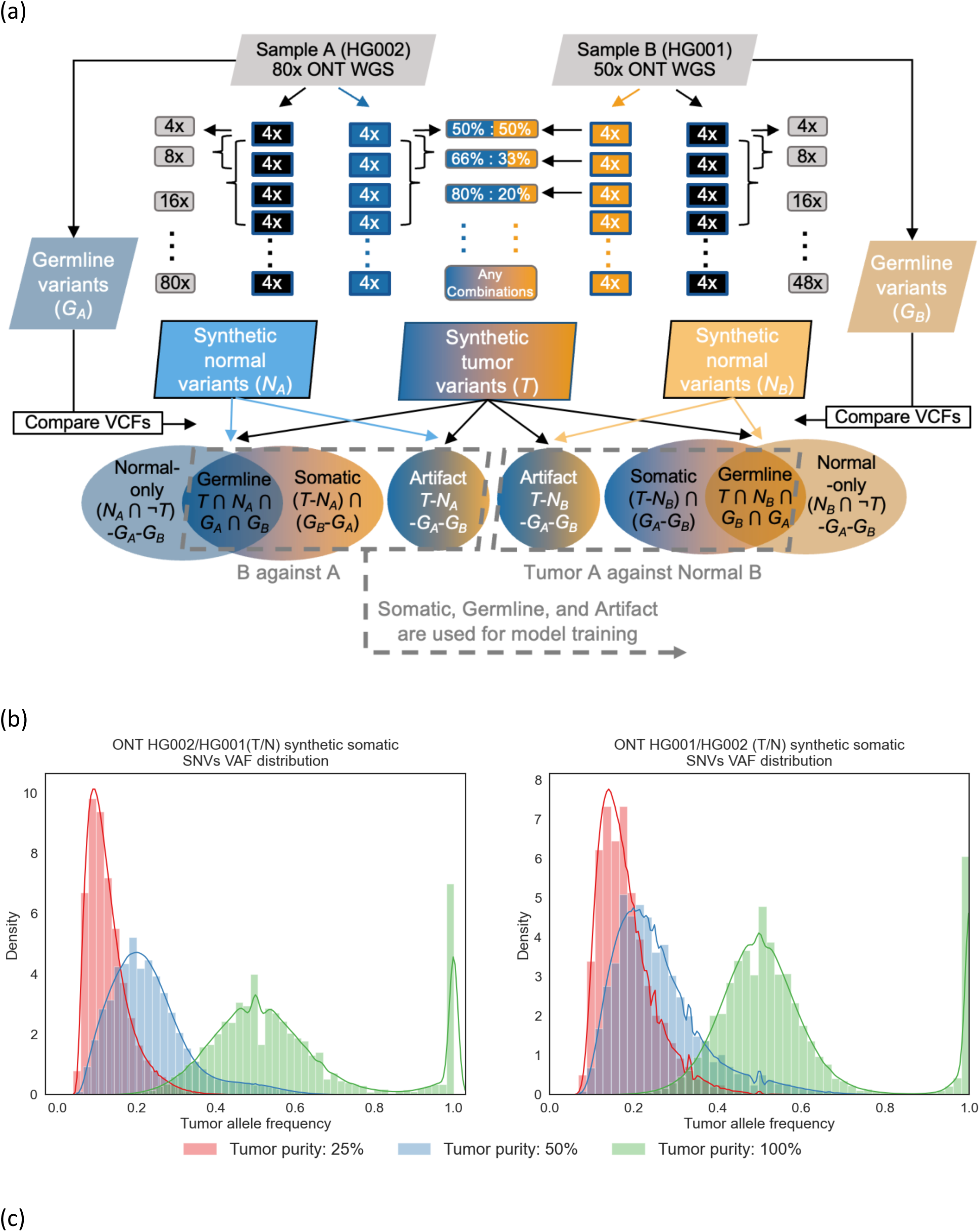
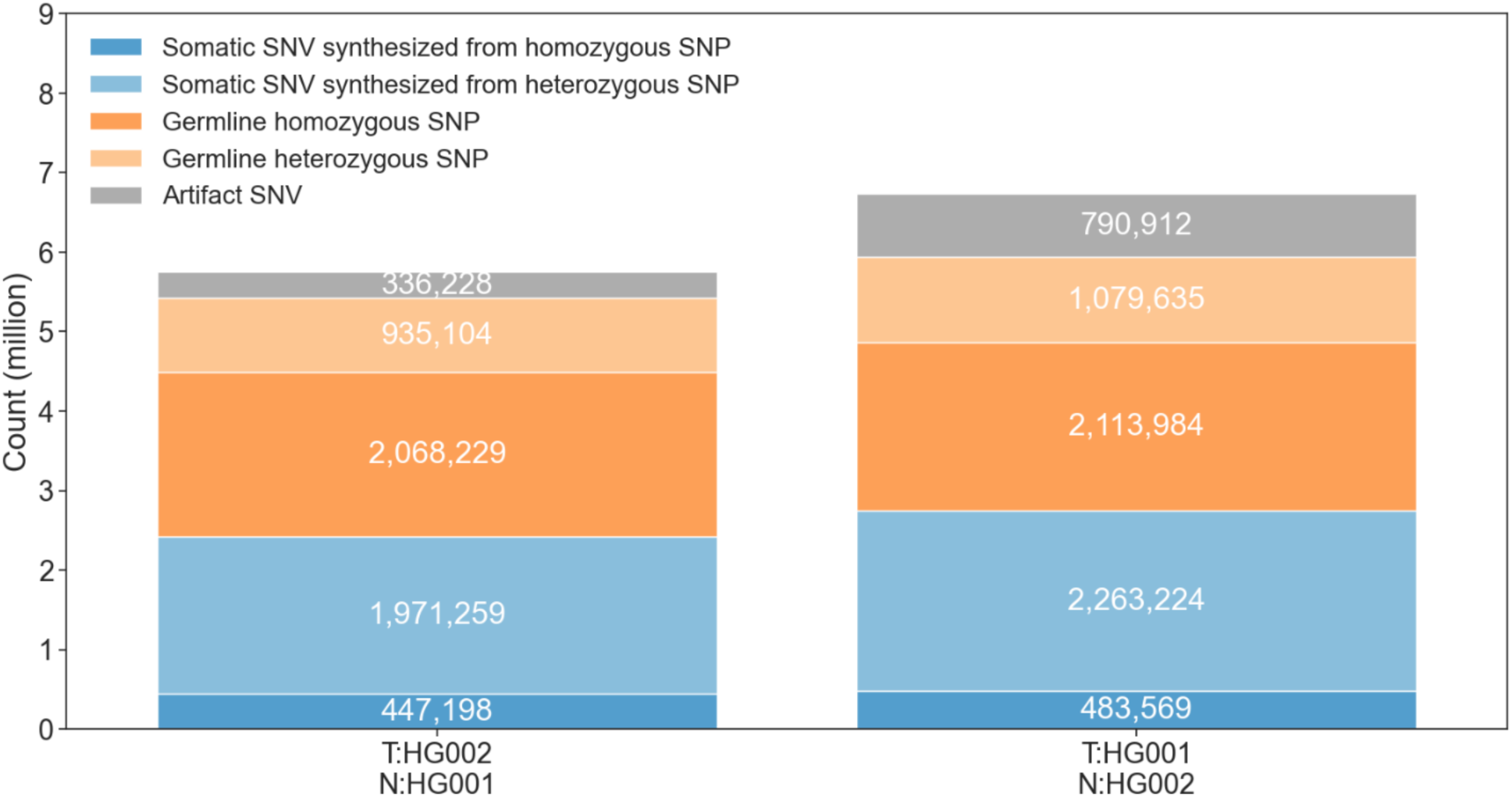
Overview of ClairS training data synthesis workflow. (a) The workflow demonstrates how to produce synthetic somatic variants using two biologically unrelated samples with known truth germline variants for ClairS model training. In this study, specifically, we used 80x ONT WGS data of GIAB HG002 as sample A, and 50x HG001 as sample B. First, germline variants *G_A_* and *G_B_* were defined as known truth germline variants in sample A and B given by GIAB. *G_A_* and *G_B_* include both homozygous and heterozygous germline variants of a sample. To generate synthetic tumor variants *T* and synthetic normal variants *N_A_*/*N_B_* for each sample, the alignments were split into smaller chunks with 4x coverage each. Then, the chunks from both samples were combined and the variants called from them were defined as *T*. With the flexibility of combining any number of chunks from both samples, *T* effectively covered variants called at different coverages and VAF. Similarly, the chunks from a sample were combined at multiple coverages for calling synthetic normal variants *N_A_* and *N_B_*. With a small number of chunks from another sample combined into a synthetic normal, *N_A_* and *N_B_* effectively covered different contamination levels. The variants *G_A_*, *G_B_*, *T*, *N_A_* and *N_B_* were then used to generate four categories of variants – “Somatic”, “Germline, “Artifact”, and “Normal-only” – with different rules. Somatic, Germline, and Artifact match the three categories in the inference task of the ClairS network architecture. The variants of the three categories were used for model training. When using sample B as tumor and A as normal, Somatic is defined as “(*T*-*N_A_*) ⋂ (*G_B_*-*G_A_*)”, i.e., variants that were 1) found in synthetic tumor *T*; 2) not found in synthetic normal *N_A_*; 3) found as a germline variant in *G_B_*; or 4) not found in *G_A_*. Germline is defined as “*T* ⋂ *N_A_* ⋂ *G_A_* ⋂ *G_B_*”, i.e., variants that were found in all *T*, *N_A_*, *G_A_*, and *G_B_*. Artifact is defined as “*T*-*N_A_*-*G_A_*-*G_B_*”, which signifies the variants found only in *T* and not in the germlines or synthetic normal. When using sample A as tumor and B as normal, the definitions remain the same except for switching the subscripts. (b) The VAF distribution of the synthetic somatic SNVs at three different simulated tumor purities (100%, 50%, and 25%), using either HG001/HG002 or HG002/HG001 as tumor/normal. Since both heterozygous and homozygous variants were used in synthesis, at 100% tumor purity, the variants were gathered at 0.5 and 1.0 VAF. The distribution showed good coverage of typical somatic SNV VAF by the synthetic SNVs. (c) The breakdown of the number of synthetic variants for training. The numbers 1) using either HG002/HG001 or HG001/HG002 as tumor/normal, and 2) of the three categories Somatic, Germline, and Artifact, as defined in subfigure a, are shown. The number of Somatic categories is further divided into those synthesized from either homozygous SNPs or heterozygous SNPs. These numbers explain why including heterozygous SNPs in the synthesis is essential to ensure a sufficient number of synthetic somatic variants for model training.

Regarding the ClairS workflow and design, Figure 2a shows an **Overview** of the ClairS workflow. Figure 2b shows **Step 1: Germline variant calling, phasing and read haplotagging**, and Figure 2c shows **Step 2. Pileup-based and full-alignment based variant calling**. For each sample, while only a fraction of genome positions has alternative allele support and among them, only a few have the potential to be called somatic variants, we used heuristics for **Selecting variant candidates**. Details of **The design of pileup input and full-alignment input** and **The design of neural networks** are shown in Figure 3. The ways in which ClairS differs from its predecessor, Clair3, are discussed in Method. Figure 2d shows **Step 3. Search for ancestral haplotype support**, which is a post-processing step that leverages more remote alignment information to search for ancestral haplotype support to the somatic variants called in step 2. To adapt to different usage scenarios, multiple **Output** options are provided.

**Figure 2.**
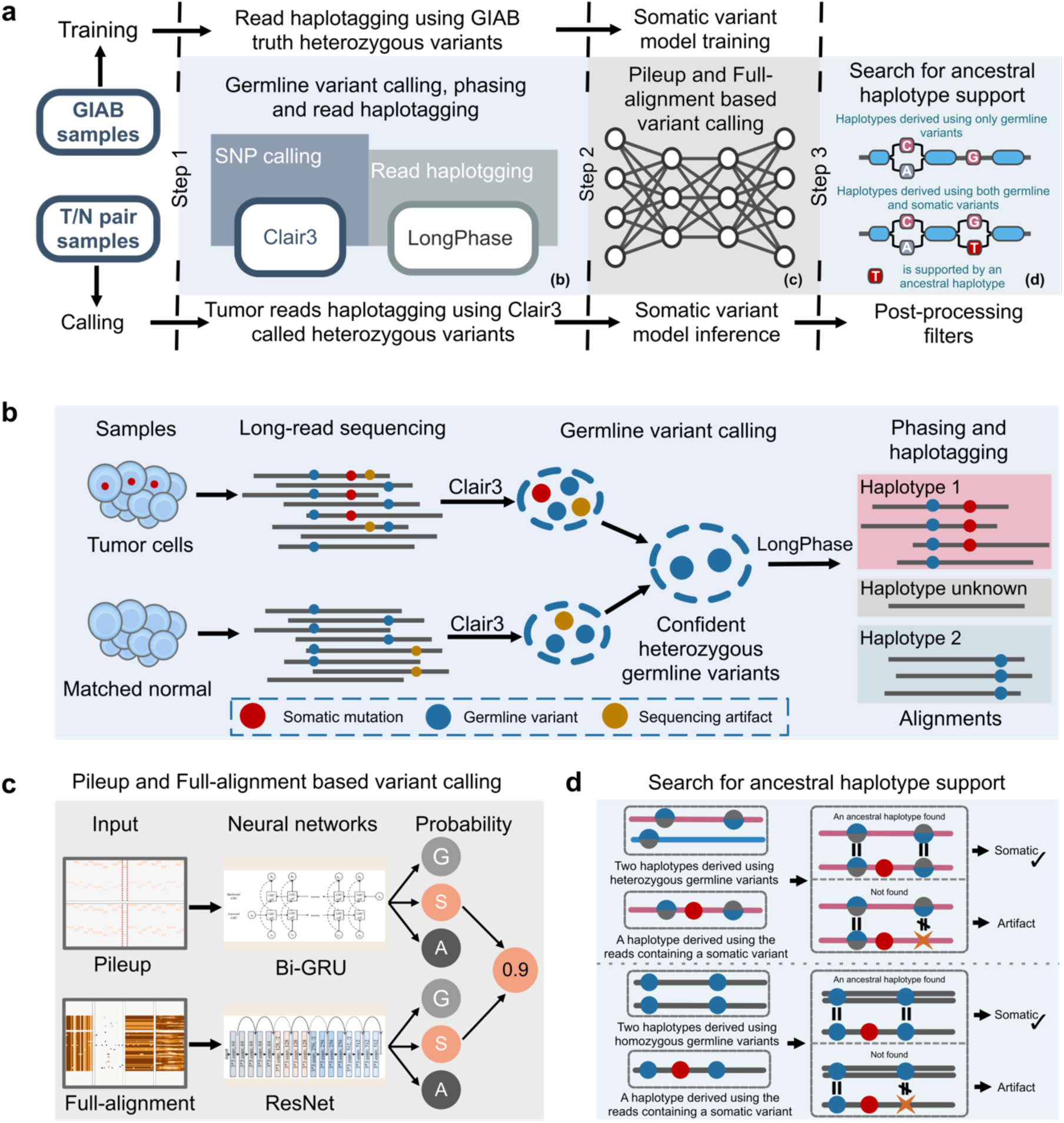
Overview of the ClairS somatic variant calling workflow. (a) The workflow illustrates the three steps of ClairS. In step 1, ClairS uses Clair3 and LongPhase for germline variant calling, phasing and read haplotagging. The processed alignments are then used for both pileup– and full-alignment-based somatic variant calling in step 2. Step 3 involves post-processing filters that eliminate somatic variant callings if an ancestral haplotype (maternal or paternal) from which the somatic variant could originate cannot be found. The details of steps 1, 2, and 3 are shown in subfigures b, c, and d. (b) Step 1 details. Clair3 is applied to both tumor and normal samples for germline variant calling. High-quality heterozygous germline variants shared by both samples are selected and used by LongPhase to phase the germline variants found in the tumor sample. Using the phased germline variants, the tumor reads are then haplotagged to belong to either haplotype 1, 2, or unknown. (c) Step 2 details. The processed alignments from step 1 are fed into both the pileup-based variant-calling neural network and the full-alignment based variant-calling neural network. On a single somatic variant candidate, both networks give respective predictions on the probability of three categories: “Somatic”, “Germline”, and “Artifact”. The predictions are then merged according to a set of rules introduced in the Method section. (d) Step 3 details. The somatic variants called in step 2 are examined to determine if they are supported by an ancestral haplotype. Ancestral haplotypes, which can be either maternal or paternal, are derived using germline variants. A somatic variant is considered supported by an ancestral haplotype if the haplotype containing the somatic variant is believed to originate from one of the ancestral haplotypes.

**Figure 3.**
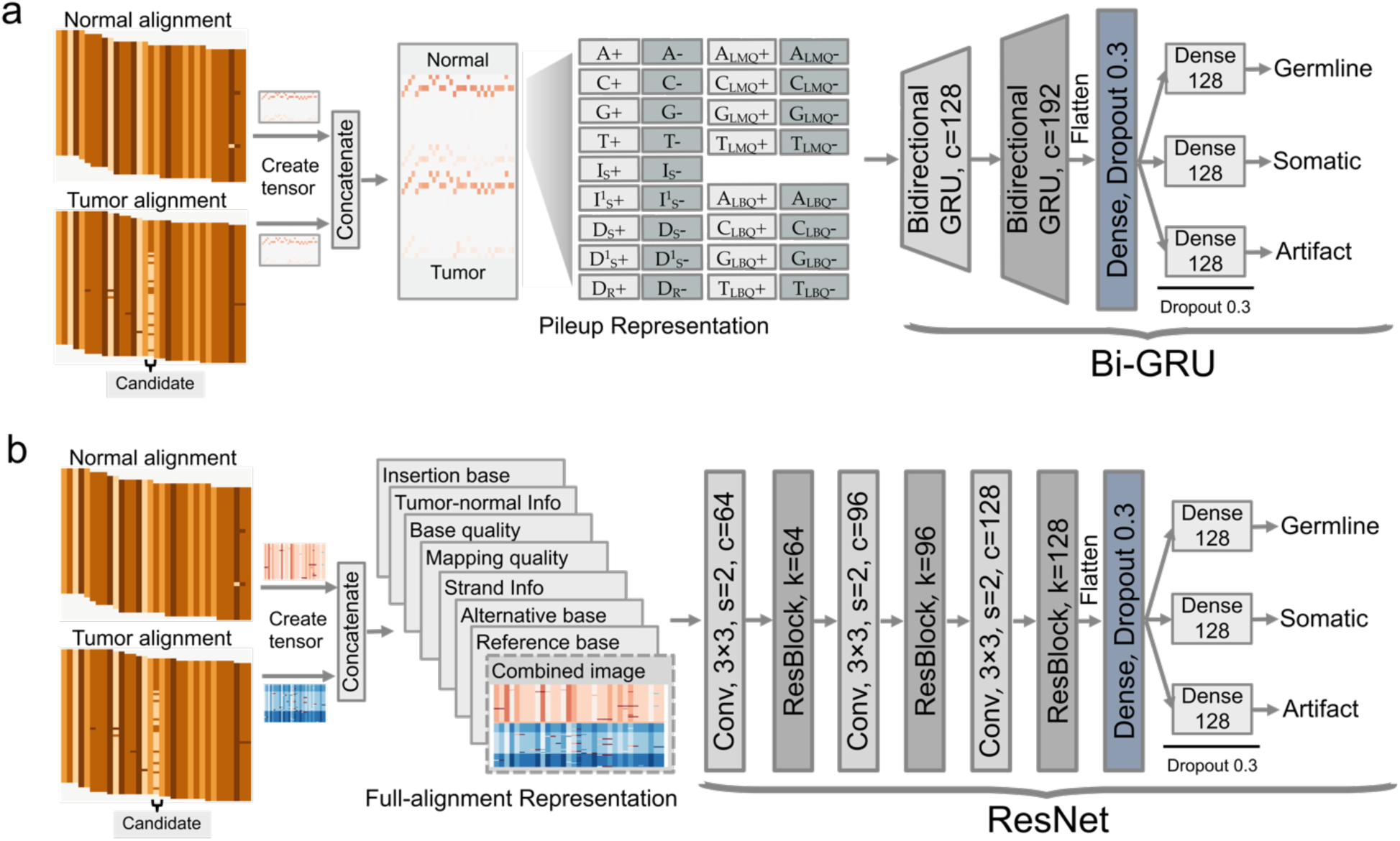
The ClairS neural network architecture. Both (a) the pileup network and (b) full-alignment network use alignments of both the tumor and normal samples as input. Tensors are created from both samples using methods detailed in the Method section, and are then concatenated. The tensors are then processed by their respective neural network for inference. Both networks output the probability of three categories: “Germline”, “Somatic”, and “Artifact”. The sequence of layers and layer configurations are shown. The letters c, s, and k, represent channel, stride, and kernel, respectively.

### ClairS performance on ONT data

A summary of the ONT data used for model training and benchmarking is shown in Supplementary Table 1. We trained the ClairS ONT model using synthetic data generated from two GIAB samples: HG001 and HG002^20^. We used both HG001/HG002 and HG002/HG001 as tumor/normal samples for data synthesis. The HG002 sample has 76.29-fold coverage and was made available by Nanopore through EPI2ME Labs. The HG001 sample has 48.44-fold coverage and was sequenced at HKU, with details given in the Method – ONT library preparation and sequencing section. Following the convention of the state-of-the-art deep-learning-based variant callers, we excluded reads and variants from Chromosome 20 from model training.

For benchmarking, we used the HCC1395/BL tumor-normal pair, with known truth variants provided by the SEQC2 consortium^3^. The HCC1395 sample has 75.97-fold coverage. The HCC1395BL sample has 45.55-fold coverage. Both samples were sequenced at two sequencing centers (HKU and Novogene) for quality control purposes. The yield of the two centers is shown in Supplementary Table 2. Only SEQC2 truth variants labeled “HighConf” (high confidence) and “MedConf” (medium confidence) were used for benchmarking. While there are ∼40k SNVs but only ∼2k Indels in the truth set, the big difference in the number of truth variants results in different analytical power between the two variant types. So we first benchmarked multiple coverages, tumor purity and normal contamination with only SNVs, and then benchmarked and discuss somatic Indel-calling performance in a separate section. We used Illumina’s Haplotype Comparison Tools^27^ to generate the performance figures, including F1-Score, Precision, and Recall, and cross-validated them with the “compare_vcf” submodule in ClairS. More clarifications and parameters are shown in the Method – Benchmarking section.

For both model training and benchmarking, we used GRCh38, which is the newest reference genome version on which both GIAB and SEQC2 truth variants are based. All ONT sequencing data were base-called using Guppy version 6.1.5 and aligned to GRCh38 using minimap2 version 2.17-r941. The command line used is given in the Supplementary Notes – Command lines used section. All data mentioned above, including ONT sequencing data, GIAB truth variants, SEQC2 truth variants, and reference genomes, are publicly accessible via links or SRA accession IDs listed in the Supplementary Notes – Data availability section.

*Performance with different tumor combinations and normal coverage.* We assessed the ClairS performance with different combinations of tumors and normal coverage. We tested three tumor coverage rates: 25-, 50-, and 75-fold. We applied 25-fold as the first step as it represents a conservative throughput estimation of an R10.4.1 PromethION flowcell. We also tested three normal coverages: 20-, 25-, and 30-fold. The 25-fold step resembles the throughput variance of a single flowcell. Our experiments aimed to imitate a practical setting for clinical cancer diagnosis, with the tumor sample coverage increased one flowcell at a time to seek higher discovery power, but the normal sample coverage is fixed at a single flowcell for cost-effectiveness.

The results are shown as Precision-Recall curves in Figure 4a. The evaluation metrics at two variant quality cutoffs (8 and 15) are shown in Supplementary Table 3. Quality cutoff 15 (hereafter referred to as “prioritize-f1 mode”) filters more variants and aims for balanced precision and recall. Quality cutoff 8 (“prioritize-recall mode”) retains more variants and aims for higher recall. In the prioritize-f1 mode, with normal coverage fixed at 25-fold, ClairS achieved 95.03%/78.71%/86.11%, 93.01%/86.86%/89.83%, and 92.94%/86.92%/89.83% precision/recall/f1-score at 25-, 50-, and 75-fold tumor coverage, respectively. From 25-to 50-fold, the recall increased from 78.11% to 86.86% (+8.75%), with a 2.02% precision drop (from 95.03% to 93.01%). From 50-to 75-fold, however, no improvement was observed. In the prioritize-recall mode, also with normal coverage fixed at 25-fold, ClairS achieved 82.66%/91.55%/86.88%, 70.21%/96.10%/81.14%, and 63.80%/96.60%/76.85% precision/recall/f1-score at 25-, 50-, and 75-fold tumor coverage, respectively. Compared to prioritize-f1, the prioritize-recall mode reached 91.55% recall at 25-fold (against 78.71%, +12.84%), and 96.10% at 50-fold (against 86.86%, +9.24%). Although Clair3 is not designed for somatic variants, to give a reference point, we conducted experiments with it using the same datasets by considering the germline variants found in tumor but not in normal as somatic variants. At 25-fold normal, Clair3 had a 19.27%/72.10%/30.42%, 29.82%/68.32%/41.52%, and 36.21%/64.56%/46.40% precision/recall/f1-score at 25-, 50-, and 75-fold tumor coverage. The results highlight the inappropriateness of using a germline variant caller for somatic variant calling.

**Figure 4.**
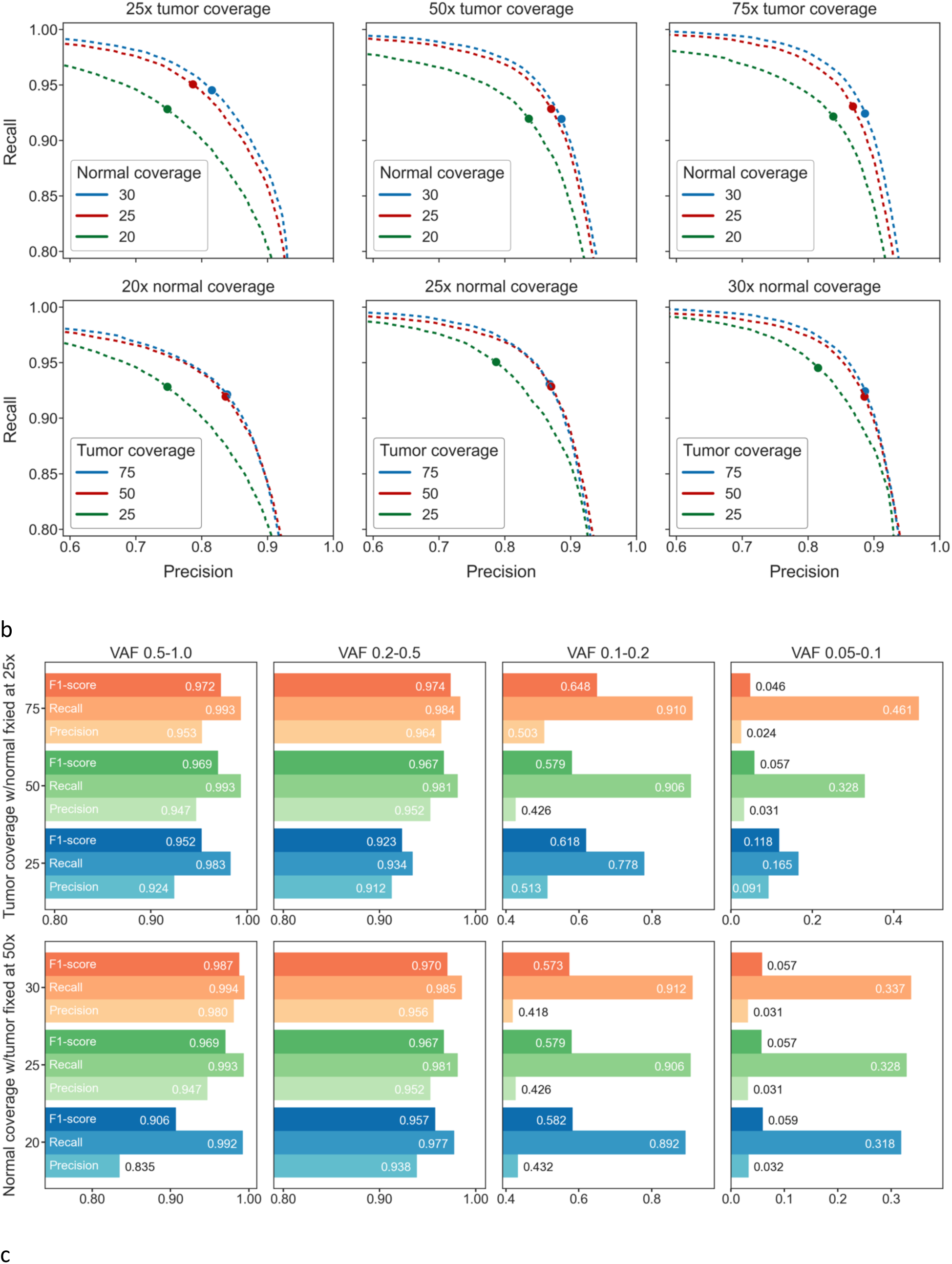
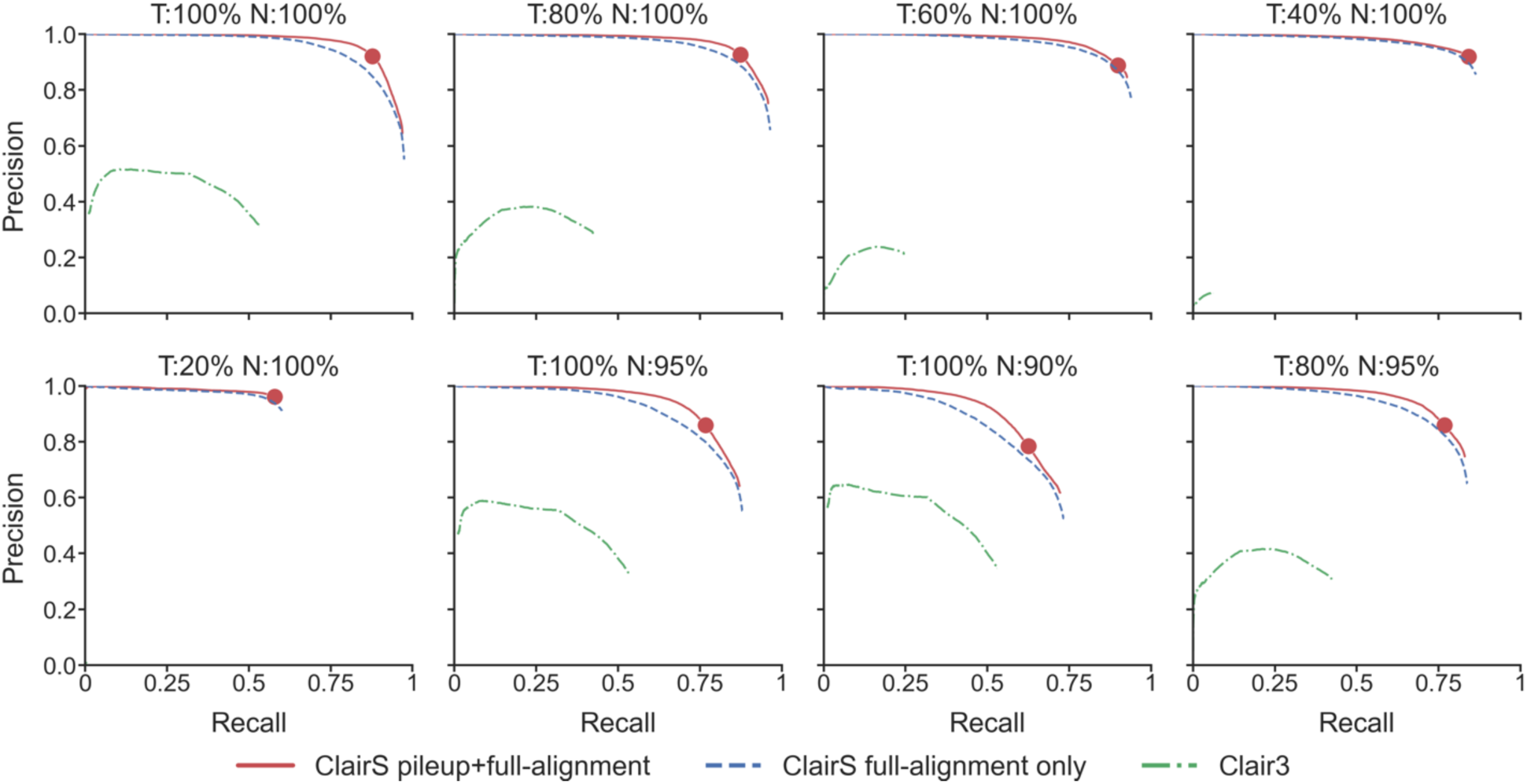
**ONT HCC1395/BL dataset benchmarking results**. (a) The precision-recall curve of different combinations of tumor and normal coverage. The dot on each dashed line shows where the best F1-score was achieved. (b) The performance of ClairS at multiple VAF ranges benchmarked on the ONT HCC1395/BL dataset. In the first row, 25, 50x, and 75x tumor were tested, with the normal coverage fixed at 25x. In the second row, 20x, 25x, and 30x of normal were tested, with tumor coverage fixed at 50x. Variant quality cutoff 8 (prioritize-recall mode) was used. (c) The precision-recall curve of different tumor/normal purity combinations with tumor coverage fixed at 50x and normal coverage fixed at 25x.

**Figure 5.**
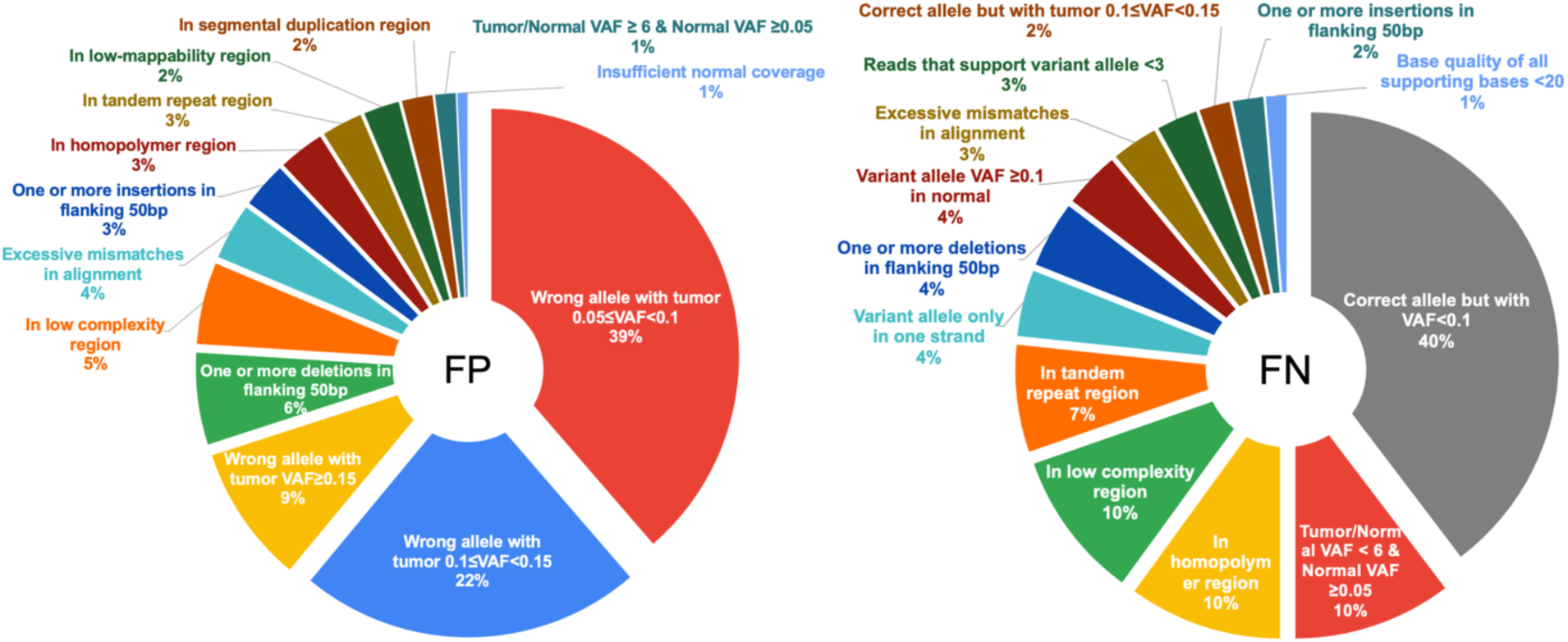
Categorizing the FPs and FNs in ClairS. The pie charts show the distribution of reasons for the FPs and FNs in ClairS. A 50x/25x ONT HCC1395/BL dataset was used for calling. 300 FPs and 300 FNs were randomly chosen from the results and analyzed. The tandem repeat, low complexity, homopolymer, and segmental duplication regions were defined using GIAB v3.0 Genome Stratification. Among the categories, “excessive mismatches in alignment” and “insufficient normal coverage” were decided manually, i.e., without certain cut-offs. “Excessive mismatches in alignment” was given if an eye check of the alignments revealed excessive inconsistent mismatches than usual alignments with a true somatic variant. “Insufficient normal coverage” was given when a germline variant signal existed in both tumor and normal, but the coverage of normal was low, so the germline variant signal in normal was obviously weaker than in tumor.

In both prioritize-f1 and prioritize-recall modes, raising the normal coverage consistently increased somatic variant-calling performance. With tumor coverage fixed at 50-fold and using the prioritize-f1 mode, ClairS achieved a 91.86%/83.77%/87.63%, 93.01%/86.86%/89.83%, and 92.06%/88.51%/90.25% precision/recall/f1-score at 20-, 25-, and 30-fold normal coverage. The numbers were 68.36%/95.69%/79.75%, 70.21%/96.10%/81.14%, and 70.56%/96.43%/81.49% in the prioritize-recall mode.

Figure 4b shows the performance of ClairS in prioritize-recall mode, broken down to four VAF ranges: 0.5-1, 0.2-0.5, 0.1-0.2, and 0.05-0.1. At different coverages, the performance of ClairS at range 0.2-0.5 (low-mid) was found to be as good as 0.5-1 (mid-high). For example, at 50/25-fold tumor/normal coverage, ClairS achieved a 94.7%/99.3%/96.9% precision/recall/f1-score at 0.5-1, and 95.2%/98.1%/96.7% at 0.2-0.5. At range 0.1-0.2, precision was reduced to ∼60%, while recalls plateaued at ∼90%. At range 0.05-0.1, precision was further reduced to below 10% (11.8%, 5.7%, and 4.6% at 25-, 50-, and 75-fold tumor coverage), while recalls were raised with increasing tumor coverage (16.5%, 32.8%, and 46.1%). The reason for the drop in precision was that the higher coverage led to a drastic increase in the number of variant candidates at very low VAF. At range 0.05-0.1, the number of candidates was about 131k, 310k, and 419k at 25-, 50-, and 75-fold tumor coverage.

*Performance at different tumor purities and normal contamination.* We assessed the performance of ClairS at different combinations of tumor purity (1.0, 0.8, 0.6, 0.4, and 0.2) and normal purity (1.0, 0.95, and 0.90). All the experiments in this section used 50-fold tumor and 25-fold normal coverage. The results are shown in Figure 4c and Supplementary Table 4. The two modes (prioritize-f1 and prioritize-recall) behaved differently with varying purity. With normal purity fixed at 1.0, in prioritize-f1 mode, precision remained above 90% (93.01%, 96.25%, 97.79%, 98.68%, and 98.99% at tumor purity 1.0, 0.8, 0.6, 0.4, and 0.2), while recall dropped (86.86%, 81.63%, 71.08%, 52.94%, and 22.43%) with decreasing tumor purity. In the prioritize-recall model, precision varied (70.21%, 80.11%, 88.14%, 93.81%, and 97.54%), while recall was boosted, especially at lower tumor purity (96.10%, 94.45%, 90.51%, 80.81%, and 53.06%). According to these results, generally, we suggest using prioritize-f1 mode at higher tumor purity and prioritize-recall mode at lower tumor purity.

Our results also showed that lower normal purity harmed somatic variant-calling performance, especially on recall. With tumor purity fixed at 1.0, in prioritize-f1 mode, the recalls were 86.86%, 70.35%, and 51.74% at normal purity 1.0, 0.95, and 0.90. In prioritize-recall mode, the recalls were 96.10%, 85.36%, and 69.42%. Accordingly, we suggest using a high-purity normal sample with ClairS for somatic variant discovery. Lastly, we showed the results using 0.8/0.95 tumor/normal purity. In prioritize-recall mode, ClairS achieved a 79.68%/81.13%/80.40% precision/recall/f1-score, demonstrating ClairS’ reliability in challenging sample conditions.

*Analysis of False Positive and False Negative calls.* Using 50-fold tumor and 25-fold normal coverage, we manually analyzed 300 false positive and 300 false negative calls randomly picked from all variant calls. Each FP and FN was assigned with the most obvious limitation as the reason why the call was false. The reasons for the 600 false calls are listed in Supplementary Table 5. A distribution of the reasons is given in Figure 4 as a pie chart.

Among the false positive calls, 39% had no matching truth but were with tumor 0.05≤VAF<0.1, 22% were with tumor 0.1≤VAF<0.15, and 9% were with tumor VAF≥0.15. As the Method section elaborates, these calls are with tumor and normal coverage ≥4, and normal VAF below 0.05. One possible explanation is that ClairS was unable to pick up more hints and tell these calls from the true ones. Some of these cases might be correctly called again with higher tumor coverage, which reduces the statistical bias in VAF. It is also possible that since the SEQC2 truth set is still under active development, its incompleteness caused a few variants that are actually true to be misclassified as false positives. Another major category of false positive calls is likely to be caused by alignment artifacts because of a repetitive or imperfect genome reference sequence. It includes 6% “One or more deletions in flanking 50bp”, 5% “In low complexity region”, 4% “Excessive mismatches in alignment”, 3% “One or more insertions in flanking 50bp”, and in total, another 10% in repetitive regions of different types. Also, we found 1% false positive calls, possibly because of insufficient normal coverage.

Among the false negative calls, 40% truth variants that were not called were with tumor VAF<0.1, 10% were with “Normal VAF ≥0.05, but tumor VAF <6 times larger than normal VAF”, and 3% were with <3 reads supporting the variant allele in a tumor. These missed variants might be called again if higher tumor coverage is given. False negative calls are more likely to be caused by alignment artifacts, as we observed 10% false negative calls in the homopolymer region, 10% in the low complexity region, 7% in the tandem repeat region, and in total, another 9% that were also likely to have been caused by alignment. We also observed 4% false negative calls caused by extreme strand bias (i.e., reads observed in only one strand); and 1% with base quality of all supporting bases <20.

*Somatic Indel-calling performance.* The SEQC2 truth set provides ∼40k SNVs but only ∼2k Indels. Owing to the scarcity of truth somatic Indels that could educe statistical biases, we benchmarked somatic Indel-calling separately. The results of different combinations of tumor coverage (25-, 50-, and 75-fold) and normal coverage (20-, 25-, and 30-fold) are shown in Supplementary Table 6. With normal coverage fixed at 25-fold, in prioritize-f1 mode (variant quality cutoff at 12) ClairS achieved 76.91%/49.36%/60.13%, 66.54%/66.89%/66.72%, and 57.79%/71.48%/63.91% of precision/recall/f1-score at 25-, 50-, and 75-fold tumor coverage. In prioritize-recall mode (cutoff at 8), ClairS achieved 53.05%/65.46%/58.61%, 38.63%/76.15%/51.25%, and 37.55%/76.60%/50.40%. As in our conclusion for SNV calling, generally, we suggest using prioritize-recall mode for calling somatic Indels at lower tumor coverage.

*Performance after adding phasing information to the input in step 2.* The reconstruction of haplotypes, also known as haplotype-resolved assembly or phasing, greatly improved the performance of long-read germline variant calling in previous practice^22, 23^. In the Method section, we elaborate how phasing information can be used to enhance somatic variant-calling performance. Using 50/25-fold HCC1395/BL and prioritize-recall mode for benchmarking, we show in Extended Figure 1b that phasable somatic variants performed better than unphasable ones, especially at lower VAF. At VAF 0.1-0.15, phasable somatic variants had a 43.8%/85.6%/57.9% precision/recall/f1-score, but unphasable somatic variants had only 16.4%/81.2%/27.3%. At VAF 0.05-0.1, phasable somatic variants had 5.8%/46.6%/10.3%, but unphasable somatic variants had only 2.3%/27.2%/4.3%. We tried disabling phasing in ClairS and fed no phasing information to the calling networks. As shown in Supplementary Table 7, the overall F1-score dropped from 81.14% to 78.66% (–2.48%).

*Performance of the two respective networks in step 2.* In contrast to Clair3, in which the pileup network handles all the variant candidates, and the full-alignment network processes only the undecided candidates using the pileup network, ClairS uses both networks equally to make collective decisions. The rationale and details are elaborated in the Method section. We tested the performance using both the pileup network only and the full-alignment network only, with 50/25-fold of HCC1395/BL and prioritize-recall mode. The results are shown in Supplementary Table 7. When we used only the pileup network, the F1-score dropped from 81.14% to 73.27% (–7.87%). When we used only the full-alignment network, the F1-score dropped from 81.14% to 79.14% (–2.00%).

*Performance of Step3: Searching for ancestral haplotype support.* As elaborated in the Method section, step 3 utilizes remote alignment signals that could not be included in the network inputs due to computational limitations to improve the precision of the called somatic variants. As shown in Supplementary Table 7, without this step, the precision dropped from 70.21% to 67.14% (–3.07%, using 50/25-fold HCC1395/BL and prioritize-recall mode).

### ClairS performance on Illumina data

Short-read somatic small-variant calling has been intensively studied. A non-exhaustive list of state-of-the-art methods include Strelka2^10^, Mutect2^8^, Lancet^6^, Neusomatic^5^, Octopus^12^, SomaticSniper^9^, and Varnet^4^. ClairS was designed primarily for long-read somatic small-variant calling. However, its poses no limitations to small reads, and we expect a variant-calling method that works for long reads to perform as well as or even better than existing short-read methods. Also, benchmarking against other short-read somatic small-variant callers provides insights on how ClairS performs against existing methods, since no other long-read callers are available for comparison.

A summary of the Illumina data used for model training and benchmarking is shown in Supplementary Table 1. We used both HG003/HG004 and HG004/HG003 as tumor/normal samples for data synthesis. The HG003 and HG004 samples had 91.15– and 88.49-fold coverage, and were publicly shared by Google Health Center. For benchmarking, we also used the HCC1395/BL tumor-normal pair, but with data from six sequencing centers, made available by the SEQC2 consortium – NS: NovaSeq at Illumina; NC: HiSeq at the National Cancer Institute, IL; HiSeq at Ilumina, EA; HiSeq at European Infrastructure for Translational Medicine, FD; HiSeq at Fudan University, NV; HiSeq at Novartis – with coverage ranging from 37.93– to 87.54-fold. The six multi-center replicates enabled us to verify ClairS’ performance consistency. If not specifically mentioned, other training and benchmarking details are the same as those of the ONT data experiments, and we benchmarked only SNVs for all callers. Like the ONT data experiments, the command lines and links to the data used for both model training and benchmarking are listed in the Supplementary Notes.

The evaluation metrics of the eight callers on the six datasets are shown in Supplementary Table 8. Figure 6a shows the precision-recall curves, and figure 6b shows a histogram of the F1-scores. ClairS consistently performed comparably or slightly better than the two top-performing callers, Strelka2 and Mutect2. On the six datasets, ClairS achieved a 97.88%, 97.82%, 97.46%, 97.53%, 97.03%, and 96.41% F1-score. Strelka2 achieved a 96.16%, 96.18%, 97.03%, 97.35%, 96.32%, and 95.47%. Mutect2 achieved 95.21%, 95.75%, 95.33%, 96.24%, 96.46%, and 94.35%. Broken down into different VAF ranges, the ClairS performance was also consistently comparable to or better than that of other callers, as shown in Figure 6c. Figure 6d shows Venn diagrams of the overlaps of false positive calls between Strelka2, Mutect2, and ClairS. The diagrams show that although rare, there are 30 to 41 false positive calls in the six datasets that were called by all three callers. Owing to the possible incompleteness of the truth set, a false positive variant can be either a wrong call or a true variant missing from the truth set. These false positive variant calls called by all three callers are worth conducting further verification and might further contribute to the completeness of the truth set.

**Figure 6.**
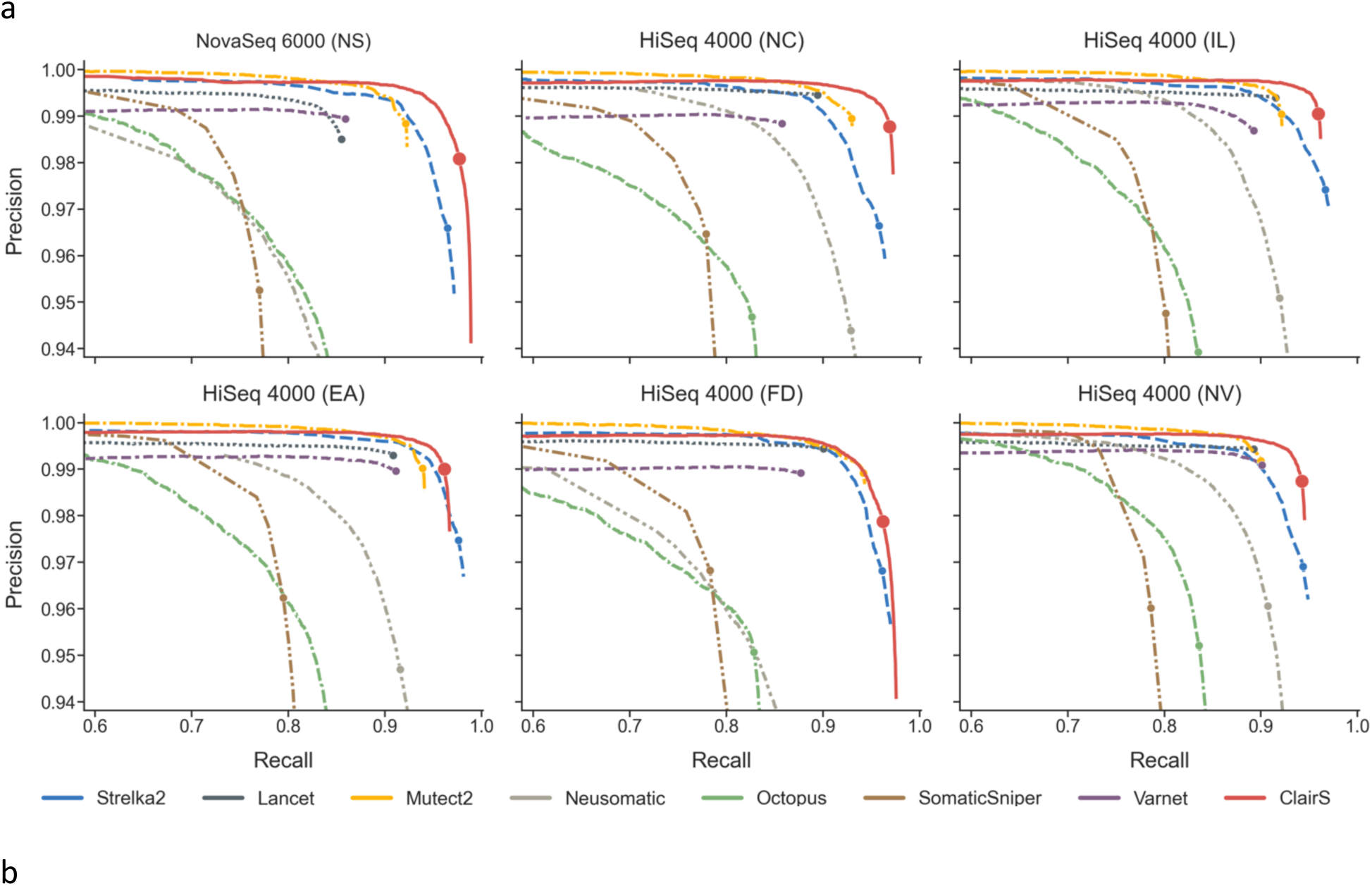
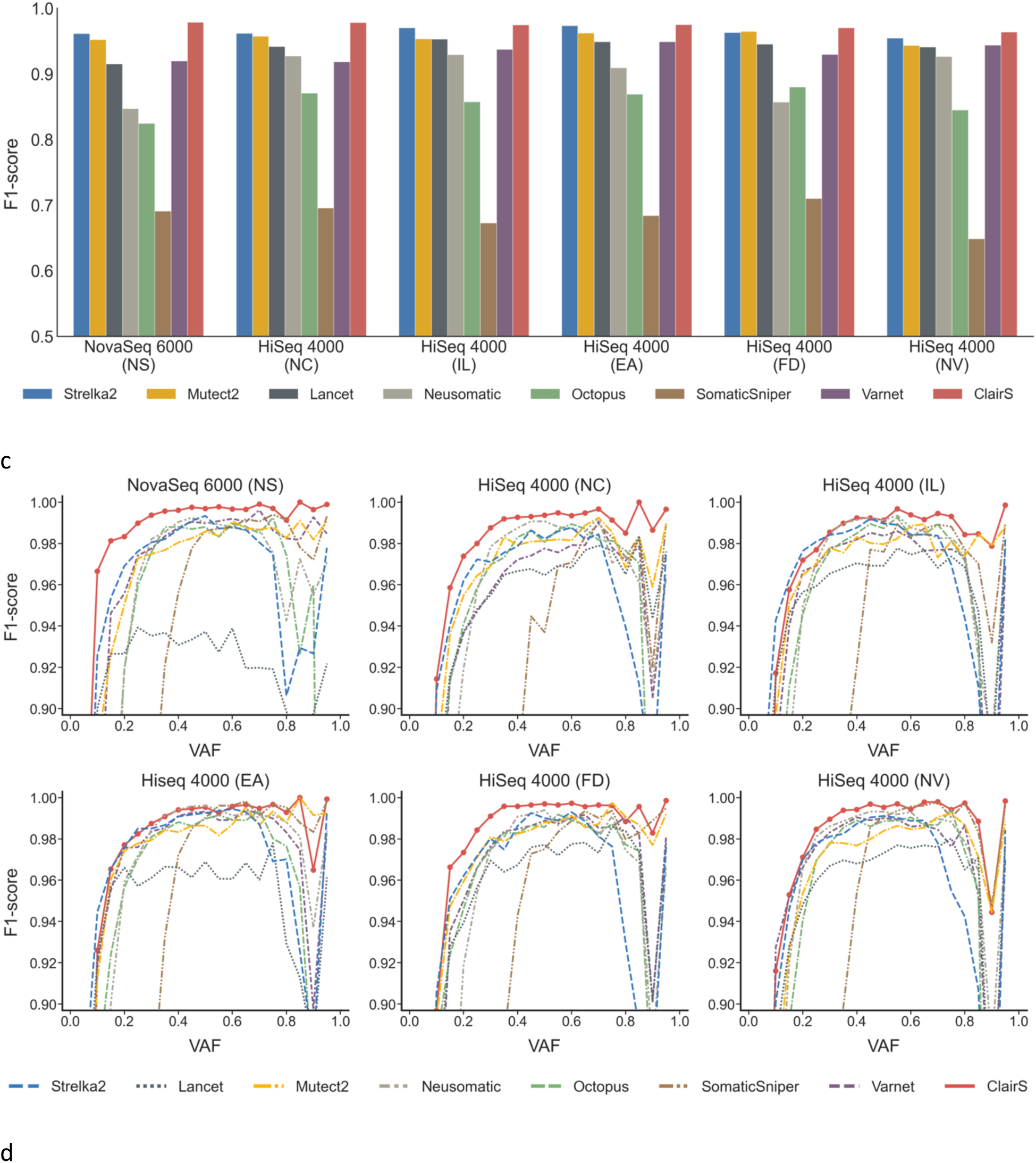
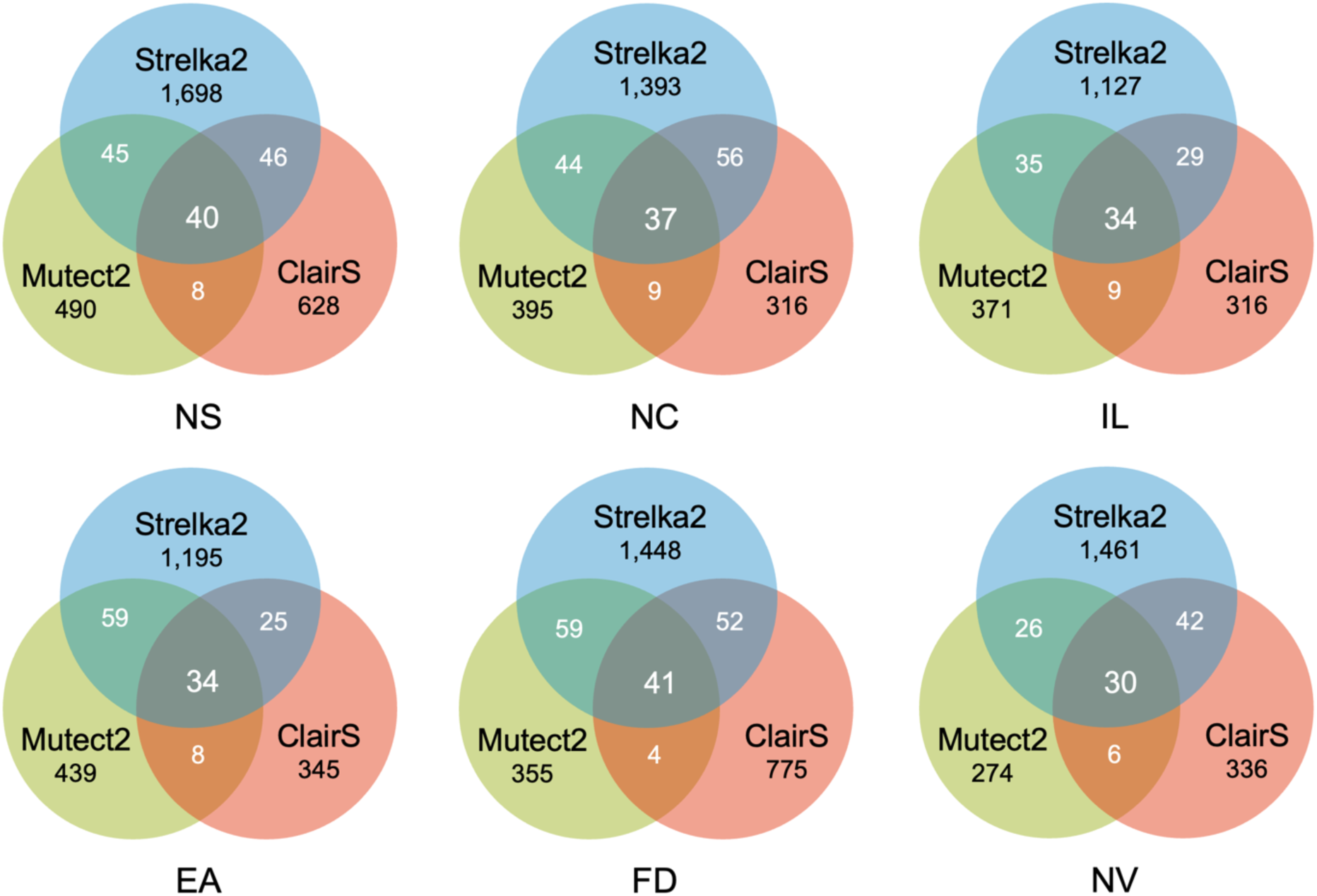
**Ilumina HCC1395/BL dataset benchmarking results**. (a) The precision-recall curve of HCC1395/BL short-read datasets from six SEQC2 sources (NS: NovaSeq at Illumina, NC: HiSeq at National Cancer Institute, IL: HiSeq at Ilumina, EA: HiSeq at European Infrastructure for Translational Medicine, FD: HiSeq at Fudan University, NV: HiSeq at Novartis) using eight tools (Strelka2, Lancet, Mutect2, Neusomatic, Octopus, SomaticSniper, Varnet, ClairS). Variants were ranked by Strelka2 – SomaticEVS, Mutect2 – TLOD, VarNet – Score, SomaticSniper – SSC, and other callers – QUAL. The dot on each line shows where the best F1-score was. (b) The overall F1-score of the experiments shown in subfigure a. (c) The F1-score at different VAFs of the experiments shown in subfigure a. (d) Venn diagrams showing the overlap of false positive variant calls between Strelka2, Mutect2, and ClairS.

## Discussion

In this study, we present ClairS, the first somatic small variant caller for ONT long-reads. In our benchmarks, we showed that it is reliable at different sample coverages, tumor purities and normal contaminations. With the training data synthesis method we devised, ClairS can be trained for somatic small-variant calling for any sequencing platform. We demonstrated that ClairS performed as well as or even better than the top-performing somatic variant callers for Illumina short reads. ClairS draws on its germline variant caller predecessors’ experience, while using a redesigned workflow, network architecture, network output, and post-processing procedure for the more challenging somatic variant-calling tasks.

The use of long reads for somatic SV discovery has unraveled complex somatic SVs that were previous hampered by short reads^28^. With the unprecedented power of long reads to cover repetitive genome regions, we expect the use of long reads for somatic small variant calling to reveal more somatic variants that were previously inaccessible by short reads, and lead to a better understanding of the mutational processes and functional consequences of the somatic variants in different cancer types. To allow more researchers to achieve these goals, ClairS was included as the small variant caller in ONT’s somatic variant-calling workflow^29^.

Despite using ONT’s latest Q20+ data, the F1-score for somatic indels was only ∼60%. Although the performance looks better when considering only the somatic indels in the coding sequences, the promise of better whole genome somatic indel-calling performance lies in the continuous advancement of ONT’s sequencing chemistry and base-calling algorithm.

## Method

### Training data synthesis

*Generating synthetic tumors and synthetic normal.* A tumor comprises normal cells and tumor cells; the latter are regarded as foreign. Similarly, the normal cells of an individual are considered foreign to the normal cells of another individual, and vice versa. The germline variants unique to an individual can mimic a somatic variant when mixed with another individual. With insufficient known truth somatic variants and standard tumor-normal sample pairs available, this observation lays the foundation for generating ample synthetic somatic variants from known truth germline variants in the GIAB reference samples with real sequencing data for deep-learning model training. The detailed workflow is shown in Figure 1a. Using 80-fold GIAB HG002 of ONT WGS alignments as the source of the tumor (hereafter referred to as A) and 50-fold of HG001 as the source of normal (B) as an example, we first split the alignments of both samples into smaller chunks, each with 4-fold coverage. The smaller chunks from both samples can be combined to simulate 1) different allele frequencies, such as combining 40-fold A, i.e., 10 4-fold chunks of A with 20-fold B, i.e., 5 4-fold chunks of B, to simulate a synthetic tumor sample with an ideal 67% allele frequency, i.e., 40-fold of A against 60-fold of A+B; 2) different coverage of both tumor and normal, e.g. increase or decrease the number of chunks as needed; and 3) different levels of contamination in normal, e.g., instead of using 100% A as normal, adding one or more chunks of B in normal.

*Coverage advice for the source data.* We applied two restrictions to tumor and normal synthesis. First, to avoid any biases caused by reusing individual reads, we used sampling without replacement in tumor and normal synthesis. That is, a chunk that was used could not be used again. Second, we required the synthetic tumor to have equal or higher coverage than the synthetic normal to align with common practice. Using the previous example, i.e., 80-fold A as the source of a tumor, and 50-fold B as the source of normal, if 20-fold B is reserved to mix with A for the tumor, then 30-fold B is left for normal, but we can achieve tumor purity only between 33% (10-fold A + 20-fold B) and 100% (80-fold A + no B). However, if 30-fold B is reserved to mix with A for the tumor, tumor purity between 0% (no A + 30-fold B) and 100% (80-fold A + no B) can be achieved, but only 20-fold B is left for normal. Thus, we suggest using the sample with higher coverage as the source of normal to achieve both higher normal coverage and full-spectrum tumor purity for model training. This is counterintuitive because higher tumor coverage of common in cancer studies.

*Deriving multiple categories of variants from a synthetic tumor/normal pair.* Four categories of variants, namely “Somatic”, “Germline”, “Artifact”, and “Normal-only”, were derived from a synthetic tumor/normal pair, as explained in detail in Figure 1a. We used GIAB truth variants in the synthetic tumor and synthetic normal, and we defined any truth variants (including both homozygous and heterozygous) as “callable-variant”. Basically, somatic variants are callable variants in the synthetic tumor but not in the synthetic normal, which are also known as truth germline variants in the tumor source, but not in the normal source. Germline variants are callable variants in both the synthetic tumor and synthetic normal, which are also known as truth germline variants in both the tumor source and normal source. Artifact variants are callable variants only in the synthetic tumor. They are not known as truth in either the tumor source or the normal source. Normal-only variants are callable-variants found only in the synthetic normal. Three of the four categories, Somatic, Germline and Artifact, are used for model training. Only the variants located in the overlapping GIAB-defined high-confidence regions of both the tumor and normal sources were used for training to ensure the quality of the training variants. Because of the sub-sampling process, some variants might have few supporting reads in the synthetic tumor, especially for low AF somatic variants. These variants were excluded from model training to avoid confusing the neural network. More exclusion details are given in the following paragraph. Our sample synthesis method supports generating synthetic tumors at any purity level, so we can use as many purities as possible to achieve fine coverage of VAF from 0 to 1, but for practicality, we used three tumor purities (25%, 50%, and 100%), and applied subsampling to all variants from the three purities to achieve acceptable VAF distribution. This is feasible because 1) the innate variance of the AF of the germline variants from the tumor and normal sources enables a pool of somatic variants fully covering VAF from 0 to 1, even with just three purities, and 2) applying subsampling to the pool enables us to enrich difficult somatic variants and reduce the number of less common somatic variants for model training. In terms of subsampling, the VAFs of chosen somatic variants were randomly selected from a beta distribution with shape parameters α=2 and β=5. The same distribution was used by the SEQC2 consortium for spike-in somatic variants in Sahraeian et al^30^. Our experiment showed that subsampling itself resulted in a ∼1.8% increase in the F1-score using 50x/25x of HCC1395/BL. The resulting VAF distribution of SNVs is shown in Figure 1b. We tried adding one more tumor purity at 12.5%, but apart from longer model training time, no performance gain was observed.

*Other details about the variants selected for model training.* For the somatic variants used for model training, a minimum coverage of four, and a minimum of three reads supporting the somatic variant allele are required. Somatic variants with VAF > 0.03 in the synthetic normal were excluded from training to avoid confusing the model with a very noisy normal. For the artifacts, the non-reference AF was capped at 0.05 to avoid using a large number of obvious artifacts for training. For germline variants, minimum coverage of four reads and a minimum of three reads supporting the germline variant allele, were required in both synthetic tumor and normal. Germline variants with a difference in AF larger than 0.1 between the synthetic tumor and normal were excluded from training. To prevent the model from inferring a somatic variant from its co-existence with two or more adjacent germline variants, which is a confounding factor that can be easily learned by the model, germlines variants that were less than 33bp (the window size of our model design) from each other were excluded from training. Our experiment showed that this exclusion alone increased somatic variant calling precision by ∼1%. With three tumor purities and the exclusions explained above, 12,489,342 training samples were left. The breakdown is shown in Figure 1c.

*Phasing information enhances somatic variant-calling performance.* An authentic somatic variant usually originates from either the maternal or paternal haplotypes, while a random error usually has a fair chance happening in both (Extended Data Figure 1a). Thus, somatic variants that have a single ancestral haplotype (either maternal or paternal) should be considered more reliable than those with two ancestral haplotypes, except for somatic variants with high VAF that might be a result of copy number alteration or clonal duplication^31^. ClairS uses phasing information for both model training and inference. Clair3 and LongPhase are used for phasing and read haplotagging. More details are given in the “ClairS input and output” section. ClairS uses phasing information during full-alignment-based variant calling, in which a channel named “Tumor/Normal/Phasing Info” is used. In this channel, the alignments are grouped into haplotype-unknown, haplotype 1, and haplotype 2, each using the read order of the alignments. Although long-read sequencing enables outstanding phasing performance, some somatic variants in difficult genomic regions or without a heterozygous germline in their vicinity still cannot be covered by any phased reads. Thus, during model training, for each variant that has a heterozygous origin from the tumor source, if one or more reads can be phased, both a version of input with reads after phasing and a version before phasing were used.

### ClairS workflow and design

*Overview.* Figure 2 shows an overview of the ClairS somatic variant-calling workflow. Starting from the alignments in the BAM/CRAM format of a tumor/normal sample pair, ClairS follows three steps to derive the somatic variants in a tumor and outputs them to a VCF file. In step 1, ClairS uses Clair3 and LongPhase for germline variant calling, phasing and read haplotagging. The processed alignments are then used for both pileup and full-alignment-based somatic variant calling in step 2. Step 3 involves post-processing filters that eliminate somatic variant calling if an ancestral haplotype (either maternal or paternal) from which the somatic variant could originate cannot be found.

*Step 1: Germline variant calling, phasing and read haplotagging.* Step 1 is depicted in Figure 2b. Clair3^22^ is integrated into ClairS for calling high-quality heterozygous germline variants in both tumor and normal to maximize the performance of the subsequent phasing task. Unlike Clair3’s default, AF≥0.2 and coverage≥10 were applied to ensure the quality of the called variants and reduce computational overhead. Only the heterozygous germline variants found in both tumor and normal were chosen for phasing. For phasing and haplotagging the tumor alignments, both LongPhase^32^ and WhatsHap^33^are allowed in ClairS. We chose LongPhase over WhatsHap as the default because LongPhase runs ∼15 times faster while delivering similar or longer phase sets on human samples. Notably, ClairS does not phase and haplotag the normal alignments. Our experiment showed phasing the normal alignments doubled the processing time but did not result in any improvement in calling performance.

*Step 2. Pileup-based and full-alignment-based variant calling.* Step 2 is depicted in Figure 2c. For a variant candidate (explained in “Selecting variant candidates”), a pileup input and a full-alignment input are generated (explained in “The design of pileup input and full-alignment input”). Then the inputs are sent to a Bi-GRU-based pileup-calling neural network and a ResNet-based full-alignment-calling neural network (explained in “The design of neural networks”) for inference. Both networks have the same output – a single task with three categories, “Somatic”, “Germline”, and “Artifact”, which match exactly the three categories defined in the synthetic training data. In contrast to Clair3, in which the faster pileup-based calling cleans up most variant candidates that are obvious variants, and the more computational-demanding full-alignment-based calling handles the tricky and less obvious candidates, ClairS considers the power of the two neural networks equal. We observed that full-alignment-based calling is performant at mid-range VAFs. However, pileup-based calling requires less evidence than full-alignment calling to draw the same conclusion. When VAF goes under 0.1, pileup-based calling becomes increasingly more sensitive and usually outperforms full-alignment-based calling. This observation makes pileup-based calling more important for somatic variant calling than its role in Clair3 for germline variant calling, especially in multiple clinical usage scenarios when sensitivity is emphasized. In ClairS, a somatic variant is called when both networks give somatic the highest probability. The variant quality (QUAL) is Phred-like and is calculated as 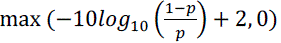, where 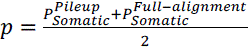. Also in contrast to Clair3, which uses the same network for both SNP and Indel calling, ClairS uses two different networks respectively trained for SNV and Indel calling. This means that ClairS runs on four networks in total: pileup for SNV, pileup for Indel, full-alignment for SNV, and full-alignment for Indel. The rationale behind the new design is that unlike germline variants that are commonly diploid, somatic variants have no ploidy assumption, meaning that the existence of SNVs and Indels in the same position are independent events. Our tests found that using separated networks led to a 1.5% increase in SNV recall. The use of separated networks also allowed the use of different variant quality cutoffs for SNV and Indel, which is useful for somatic variant calling, especially when the sample condition is not ideal.

*Selecting variant candidates.* Sending every genome position as a variant candidate to the neural networks guarantees maximum sensitivity. However, it is not only computationally infeasible, but also unreasonable to work on nonstarter positions, such as those without any non-reference allele support. A good variant candidate selection strategy is essential to achieve a balance between sensitivity and running time. In ClairS, the selection criteria are as follows. Let *r*∈*K*=(*A, C, G, T*) be the reference base of a genome position, and *m*∈ *K-r* be the alternative bases. *D^X^_m_* denotes the coverage of *m* at the position in sample *X*∈{*T, N*}, where *T* and *N* represent the tumor and normal sample. *C_m_* defines the selection criteria of each alternative base in *m*, as:

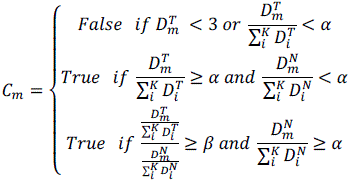

where α sets the minimum VAF, and β sets the minimum tumor VAF to normal VAF ratio for a candidate to be selected. Intuitively, the first equation means disregarding variant candidates with < 3 reads in tumor supporting the variant allele, or with VAF in tumor < α. The second equation means selecting a variant candidate if its VAF is <α in normal, but ≥α in tumor. The third equation means selecting a variant candidate, even if its VAF in normal is ≥α, the VAF in tumor is ≥β times larger than the VAF in normal. In ClairS, α and β are configurable and default to 0.05 and 6, respectively. Like model training data preparation, coverage ≥4 is required in both tumor and normal for a candidate to be selected for variant calling.

*The design of pileup input and full-alignment input.* ClairS pileup input comprises 1,122 integers – 33 positions wide with 34 features at each position (an example is given in Extended Data Figure 2a). A detailed explanation of each feature is given in Supplementary Methods under “Description of pileup input features”. ClairS has 18 pileup features in common with Clair3, and 16 additional features. The 16 new features are read counts *N_LMQ_+*, *N_LMQ_-*, *N_LBQ_+*, and *N_LBQ_-*, where *N* is either of the nucleotides A, C, G, and T, LMQ subscript means mapping quality lower than 20 (MQ<20), LBQ means base quality lower than 30 (BQ<30), and + and – mean the forward and reverse strand, respectively. The rationale behind the new features is that in ClairS, the results of pileup-based calling and full-alignment-based calling are trusted equally, so the mapping quality and base quality information that used to be exclusive to full-alignment calling need to be added to pileup-based calling. Our experiment showed that removing the 16 new features reduced precision by ∼2% using 50x/25x of HCC1395/BL. ClairS full-alignment input comprises 30,030 integers – seven channels, each with 33 positions and 130 rows to support at most 76 tumor reads, 52 normal reads, and 2 empty rows as space between tumor and normal (an example is given in Extended Data Figure 2b). Like Clair3, random subsampling down to the maximum supported coverage is used at excessive coverages. A detailed explanation of each channel is given in Supplementary Methods under “Description of full-alignment input channels”. In both inputs, the candidate variant is centered at the 16^th^ position. Positions uncovered by any base in full-alignment input are filled with zero.

*Design of neural networks.* The pileup and full-alignment network architecture and important parameters are shown in Figure 3. The pileup network uses two bidirectional gate recurrent unit (Bi-GRU) layers, each with 128 and 192 units. Compared to the Clair3 pileup network, the use of Bi-GRU instead of bidirectional long short-term memory (Bi-LSTM) architecture reduced trainable parameters from 2,532,995 to 2,309,507 and matrix computations from 3.11 to 2.38 billion, but improved performance in our experiment. The full-alignment network is a residual neural network (ResNet) comprising three standard residual blocks. A convolutional layer is added immediately before each residual block to expand the number of channels. In both networks, a dropout rate at 0.3 is set for the flattened layer and dense layer to prevent overfitting.

*Step 3. Search for ancestral haplotype support.* Step 3 is depicted in Figure 2d. The neural networks exhibited good power in distinguishing real variants from false positive candidates. However, useful signals remote to a variant candidate are not covered by the current neural network designs in ClairS, which considers only the flanking 16bp of a candidate. Notably, even if the flanking window is extended to 50bp, it is still too short for an accurate inference of which haplotype a variant candidate belongs to using only the networks, but the networks would already be computationally infeasible for somatic variant calling. In ClairS, post-processing step 3 is designed to reduce false positive calling mistakes made by the networks by leveraging relatively remote germline variants to find the correct ancestral haplotype for a somatic variant. Any somatic variant calls that cannot be found with ancestral haplotype support are switched to an artifact and are excluded from the output. The haplotagged reads produced in step 1 are used in this step. For a somatic variant that covers any haplotagged reads, we required the somatic variant to coexist with the heterozygous germline variants less than 100 bp away on its left and right in the reads in the haplotype group the somatic variant supporting reads were in. An example of a false positive somatic variant filtered by this rule is given in Extended Data Figure 3a. A somatic variant at chr4:38,012,942 was called by the two networks. A phased heterozygous germline variant was found 61 bp left of the somatic call. Three reads in haplotype 2 that supported the somatic variant were found not to have the heterozygous germline variant. Thus, the somatic variant was considered unsupported by an ancestral haplotype. For a somatic variant that covers no haplotagged read, it is probably because there are no germline variants or only homozygous variants in the vicinity. In this case, we required the somatic variant to be coexisting with the homozygous germline variants less than 100 bp away on its left and right in all somatic variant supporting reads. An example of this is given in Extended Data Figure 3b. A somatic variant at chr1:100,632,158 was called by the two networks. A homozygous germline variant was found 39 bp left of the somatic call. Multiple reads that support the somatic variant were found not to have the homozygous germline variant. Thus, the somatic variant was not considered to be supported by an ancestral haplotype. Somatic variants that do not have any germline variants less than 100 bp away on their left or right are not applicable in this step.

*Output.* ClairS supports VCF format output. Somatic variants are marked “PASS” or “LowQual” if the variant quality is low (i.e., QUAL<8, configurable by option), or they are filtered in step 3. For each variant, the allele frequency and supporting coverage of the reference allele and all alternative alleles are shown. The options “--print_germline_calls” and “--print_ref_calls” enable outputting germline variants and artifacts, respectively.

### ONT library preparation and sequencing

Genomic DNA (gDNA) of a triple-negative breast cancer (TNBC) cell line (HCC1395) and a B lymphocyte-derived normal cell line (HCC1395BL) from the same donor were purchased from the American Type Culture Collection (ATCC). Genomic DNA (gDNA) of HG001 was purchased from the Coriell Institute. The high-molecular-weight gDNA was examined by Nanodrop, Qubit, and 0.35% agarose electrophoresis for its concentration, purity, and integrity. The gDNA was then fragmented with gTube to generate DNA fragments approximately 20 kb in length. These fragments were then being sequenced at two sequencing centers: HKU and Novogene. At HKU, the fragments of HCC1395, HCC1395BL, and HG001 were prepared and ligated with a sequencing adapter using ONT’s ligation sequencing kit V14 SQK-LSK114. The ligated samples were sequenced on R10.4.1 PromethION flowcells using a PromethION 2 Solo device and MinKNOW software version 1.18.02, for 96 h. At Novogene, the fragments of HCC1395 and HCC1395BL were prepared and ligated with a sequencing adapter using ONT’s ligation sequencing kit V12 SQK-LSK112. The ligated samples were sequenced on R10.4 PromethION flowcells using PromethION 48, for 96 h.

### Benchmarking

We used the truth set of somatic variants in HCC1395/BL generated and maintained by the SEQC2 consortium. The truth set was orthogonally validated with multiple sequencing replicates from multiple sequencing centers that comprise over 1,500-fold sequencing data in total. We used only the somatic variants labeled “HighConf” (High Confidence) or “MedConf” (Medium Confidence) as truth. Somatic variants labeled “LowConf” (Low Confidence, VAF ≤0.05, not a part of the truth set as defined by SEQC2) were not used for benchmarking. In total, there were 39,560 truth SNVs and 1,922 truth Indels; 39,447 of the SNVs and 1,602 of the Indels were within the high-confidence regions defined in a BED file provided by SEQC2. A variant call was considered correct only if it matched both the genome position and variant allele of the truth. For both the ONT and Illumina benchmarks, some truth variants were excluded for the following reasons. First, even with the high sequencing coverage, such as 75.97-fold HCC1395 we generated for the ONT benchmarks, some truth variants still had very low or no coverage, or had no read supporting the variant allele. These truth variants would fail all the benchmarks, so they should be excluded. Second, some benchmarks tested multiple sequencing coverages and required sequencing read subsampling from the full dataset. The subsampling process might remove reads supporting a truth variant to an extent that few or no supporting reads are left.

This affects especially the somatic variants that already have a low VAF. For example, a VAF 0.05 somatic variant with 20-fold coverage and one read supporting the variant allele can be reduced to VAF 0 by removing just one read during subsampling. This reduces the quality of the benchmarking results, especially for low VAF truth variants when subsampled datasets are used. To alleviate the problem, any truth variants that have very low VAF (<0.05) observed in the full dataset before subsampling should be excluded. Summing up the two reasons above, for each of the full datasets we used in both the ONT and Illumina benchmarks, we excluded truth variants that matched any of the following criteria from benchmarking: 1) VAF ≤0.05, 2) reads supporting the variant allele <3, 3) tumor coverage <4, and 4) normal coverage <4. For standardization, we used som.py, provided in Illumina’s Haplotype Comparison Tools^27^ (version v0.3.12) to generate evaluation metrics, including F1-Score, Precision, and Recall against the truth variants. The “compare_vcf” submodule in ClairS produces identical results to som.py, but automates the exclusion of unqualifying truth variants. The truth set materials are publicly available to the community. All tools, their version, and command lines used are given in the “Command lines used” section in Supplementary Notes.

### Computational performance

ClairS was written in Python and C++. The Python parts leveraged PyPy for speed up. The neural network implementations used PyTorch. Training ClairS neural networks requires a high-end GPU, but using ClairS for somatic variant calling requires only a CPU. For the 50x/25x HCC1395/BL pair, ClairS finished running in ∼5 hours for ONT data and ∼2 hours for Illumina data (30% slower than Strelka2, but faster than all other short-read somatic variant callers), using two 12-core Intel Xeon Silver 4116 processors. The memory footprint is low and is controlled at lower than 1GB per CPU. For model training, we tested Nvidia GeForce RTX 2080 Ti, 3090, and 4090, and found each new model provided a ∼35% speed increase from the previous generation.

## Code availability

ClairS is open source and available at https://github.com/HKU-BAL/ClairS under the BSD 3-Clause license. The results in this paper were based on the ClairS initial release (version 0.0.1). Multiple installation options are available for ClairS, including Docker and Singularity. ClairS has also been included as the small variant caller in ONT’s somatic variant calling workflow^29^ since version 0.1.0.

## Data availability

The links to the reference genomes, truth somatic variants, benchmarking materials, ONT, and Illumina data are given in the “Data availability” section in Supplementary Notes. All analysis output, including the VCFs and running logs, is available at http://www.bio8.cs.hku.hk/clairs/analysis_result. The HCC1395/BL sequencing data generated in this study was deposited in the NCBI short-read archive with accession ID PRJNA986292.

## Acknowledgements

R.L. was supported by Hong Kong Research Grants Council grants GRF (17113721) and TRS (T21-705/20-N), the Shenzhen Municipal Government General Program (JCYJ20210324134405015), the URC fund from HKU, and Oxford Nanopore Technologies.

## Author contributions

R.L. conceived the study. Z.Z. and R.L. designed the algorithms, implemented ClairS, and wrote the paper. Y.L. and T.-W.L. evaluated the benchmarking results. All authors revised the manuscript.

## Extended Data Figures

**Extended Data Figure 1.**
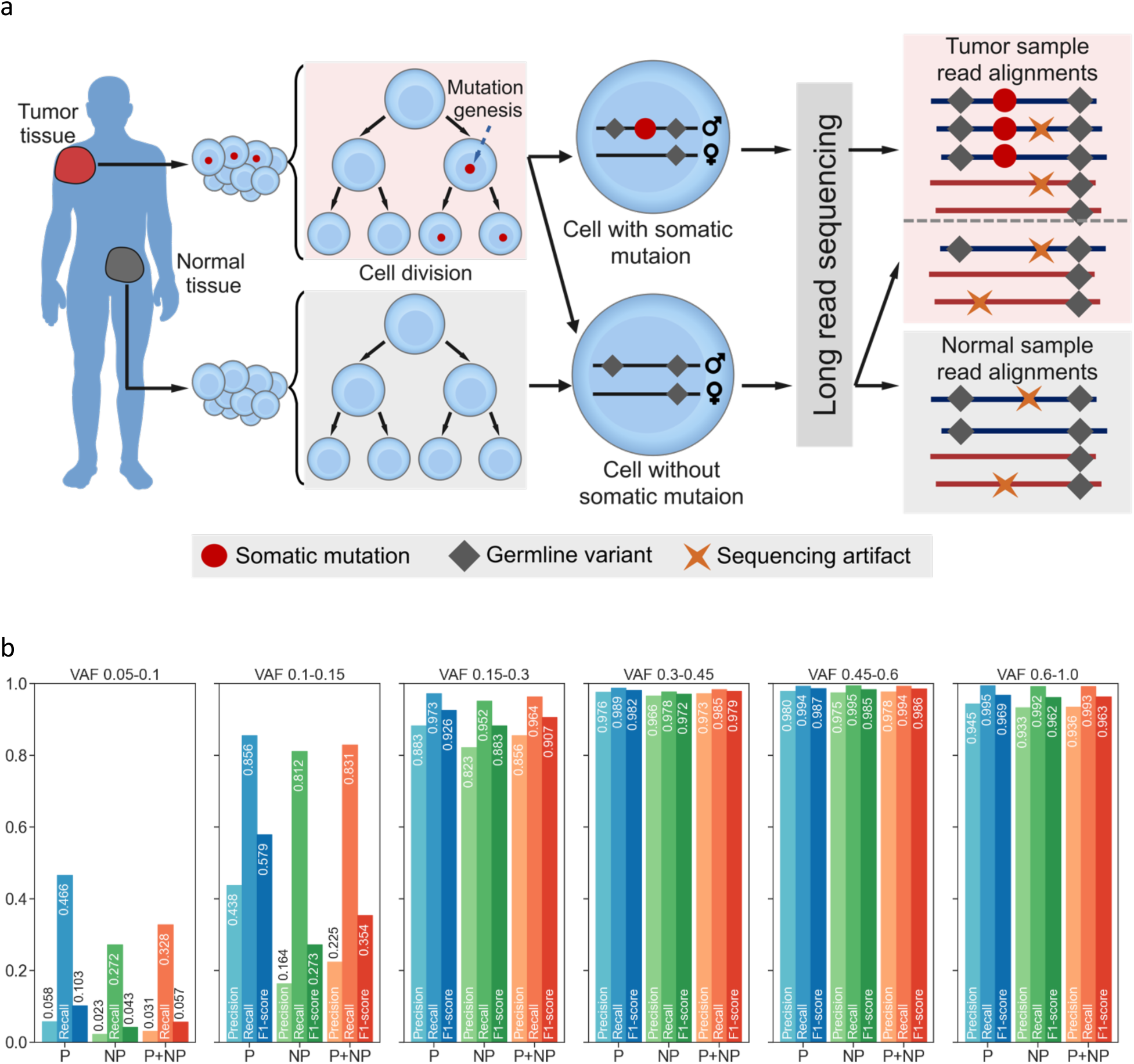
Performance differences between phasable and not-phasable SNVs. (a) The figure shows that a somatic variant usually originates in a single somatic cell and then spreads to more cells through cell division, resulting in a clonal carrying the same variant. It also shows how the mismatches in the tumor sample and normal sample are different from each other. A somatic variant is more likely to be assigned to a haplotype through phasing, while a variant caused by random sequencing errors is less likely to be successfully phased. (b) A performance comparison of somatic variants where “P”: can be phased, and “NP”: cannot. The figure shows a higher performance in somatic variants that can be phased, especially at lower VAFs. We used 50/25-fold HCC1395/BL and prioritize-recall mode.

**Extended Data Figure 2.**
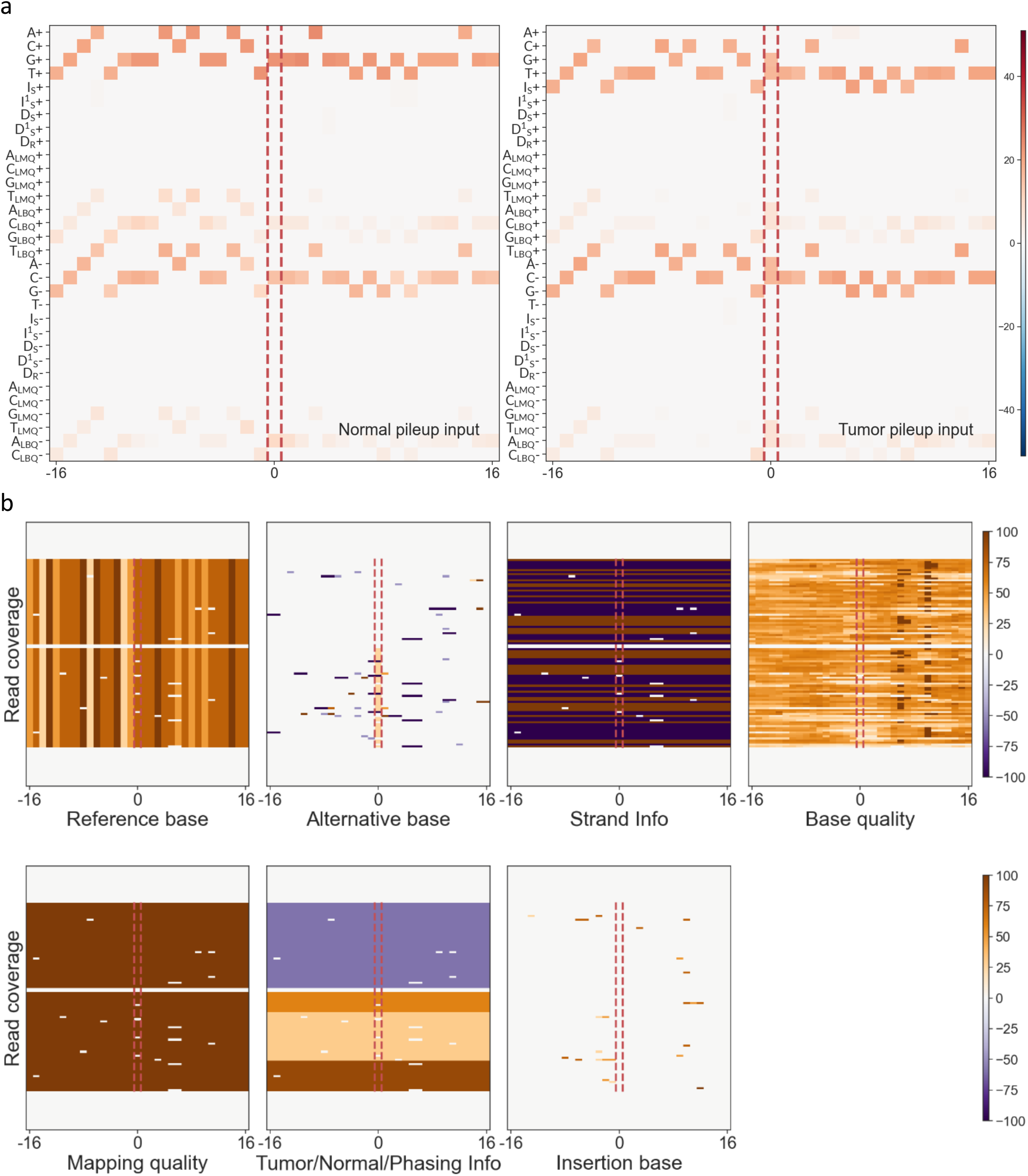
Visualization of neural network inputs. (a) Pileup-based calling input visualization. The candidate site is centered and marked by two dashed lines. (b) Full-alignment-based calling input visualization. In b, the top and bottom are padded with zero when the total coverage of tumor and normal samples does not reach the input limit. The normal read alignments and tumor read alignments in all channels are separated by two rows filled with zeros. The two demonstrations involved truth variants randomly picked from the HCC1395/BL dataset.

**Extended Data Figure 3.**
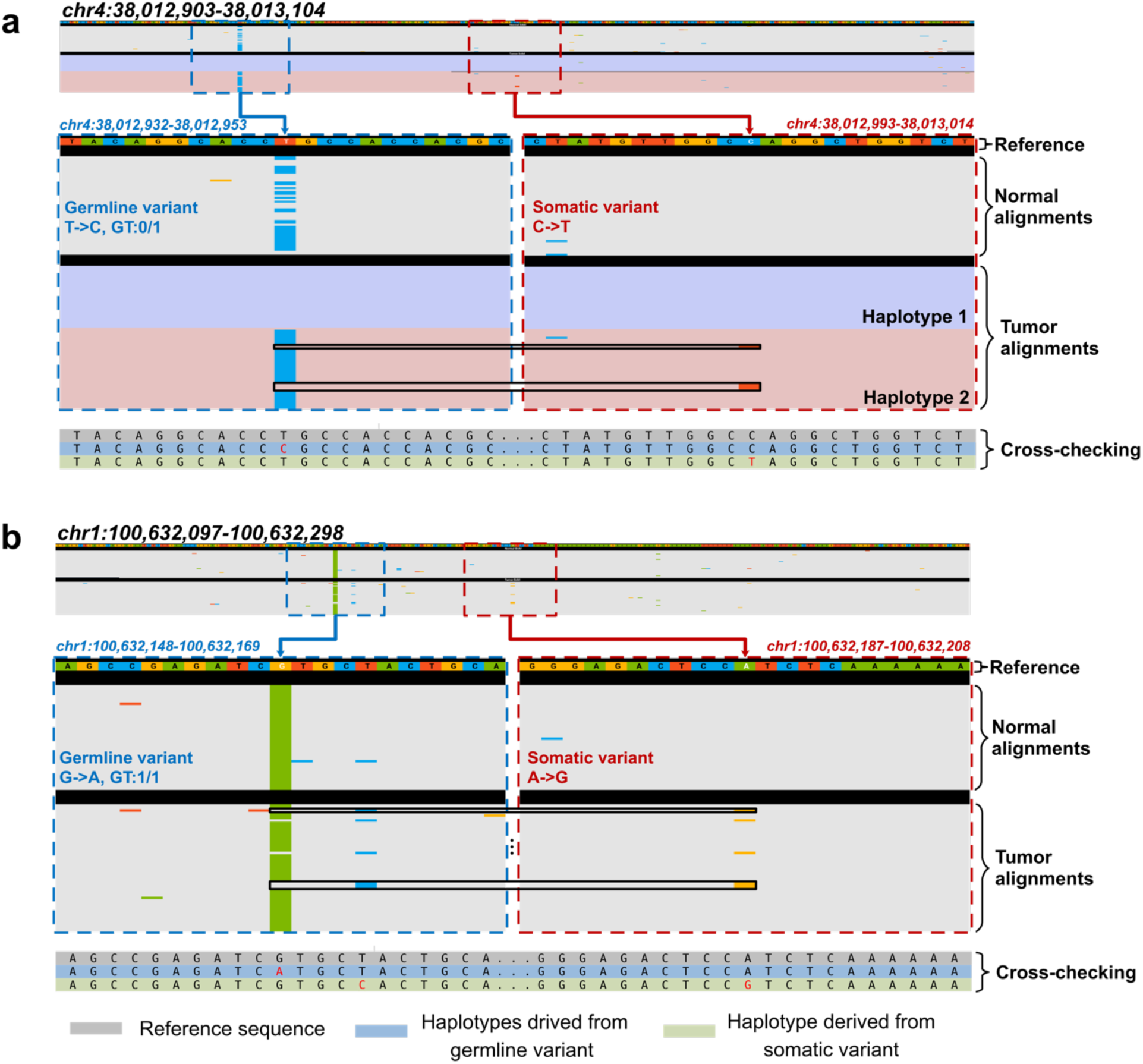
Two examples of haplotype inconsistency that signifies a false somatic call. (a) Example of a false somatic call with a haplotype inconsistent with the haplotype derived from a heterozygous germline variant nearby. (b) Example of a false somatic call with a haplotype inconsistent with the haplotypes derived from a homozygous germline variant nearby. The bases A, C, G, and T are depicted in green, blue, yellow, and red, respectively. The background in gray, purple, and pink represents an unknown haplotype, haplotype 1, and haplotype 2, respectively. GT is an abbreviation of genotype.

## References

1. Weinstein, J.N. et al. The cancer genome atlas pan-cancer analysis project. Nature genetics 45, 1113–1120 (2013).

2. Perera-Bel, J. et al. From somatic variants towards precision oncology: evidence-driven reporting of treatment options in molecular tumor boards. Genome medicine 10, 1–15 (2018).

3. Fang, L.T. et al. Establishing community reference samples, data and call sets for benchmarking cancer mutation detection using whole-genome sequencing. Nature biotechnology 39, 1151–1160 (2021).

4. Krishnamachari, K. et al. Accurate somatic variant detection using weakly supervised deep learning. Nature Communications 13, 4248 (2022).

5. Sahraeian, S.M.E. et al. Deep convolutional neural networks for accurate somatic mutation detection. Nature communications 10, 1041 (2019).

6. Narzisi, G. et al. Genome-wide somatic variant calling using localized colored de Bruijn graphs. Communications biology 1, 20 (2018).

7. Fan, Y. et al. MuSE: accounting for tumor heterogeneity using a sample-specific error model improves sensitivity and specificity in mutation calling from sequencing data. Genome biology 17, 1–11 (2016).

8. Cibulskis, K. et al. Sensitive detection of somatic point mutations in impure and heterogeneous cancer samples. Nature biotechnology 31, 213–219 (2013).

9. Larson, D.E. et al. SomaticSniper: identification of somatic point mutations in whole genome sequencing data. Bioinformatics 28, 311–317 (2012).

10. Kim, S. et al. Strelka2: fast and accurate calling of germline and somatic variants. Nature methods 15, 591–594 (2018).

11. Freed, D., Pan, R. & Aldana, R. TNscope: accurate detection of somatic mutations with haplotype-based variant candidate detection and machine learning filtering. biorxiv, 250647 (2018).

12. Cooke, D.P., Wedge, D.C. & Lunter, G. A unified haplotype-based method for accurate and comprehensive variant calling. Nature biotechnology 39, 885–892 (2021).

13. Lai, Z. et al. VarDict: a novel and versatile variant caller for next-generation sequencing in cancer research. Nucleic acids research 44, e108–e108 (2016).

14. Kovaka, S., Ou, S., Jenike, K.M. & Schatz, M.C. Approaching complete genomes, transcriptomes and epi-omes with accurate long-read sequencing. Nature Methods 20, 12–16 (2023).

15. Ameur, A., Kloosterman, W.P. & Hestand, M.S. Single-molecule sequencing: towards clinical applications. Trends in biotechnology 37, 72–85 (2019).

16. Nanopore Q20+ chemistry, https://nanoporetech.com/q20plus-chemistry. (2019).

17. Fox, E.J., Reid-Bayliss, K.S., Emond, M.J. & Loeb, L.A. Accuracy of next generation sequencing platforms. Next generation, sequencing & applications 1 (2014).

18. Luo, R., Sedlazeck, F.J., Lam, T.-W. & Schatz, M.C. A multi-task convolutional deep neural network for variant calling in single molecule sequencing. Nature communications 10, 998 (2019).

19. Poplin, R. et al. A universal SNP and small-indel variant caller using deep neural networks. Nature biotechnology 36, 983–987 (2018).

20. Wagner, J. et al. Benchmarking challenging small variants with linked and long reads. Cell Genomics 2, 100128 (2022).

21. Luo, R. et al. Exploring the limit of using a deep neural network on pileup data for germline variant calling. Nature Machine Intelligence 2, 220–227 (2020).

22. Zheng, Z. et al. Symphonizing pileup and full-alignment for deep learning-based long-read variant calling. Nature Computational Science 2, 797–803 (2022).

23. Shafin, K. et al. Haplotype-aware variant calling with PEPPER-Margin-DeepVariant enables high accuracy in nanopore long-reads. Nature methods 18, 1322–1332 (2021).

24. Smolka, M. et al. Comprehensive structural variant detection: from mosaic to population-level. BioRxiv, 2022.2004. 2004.487055 (2022).

25. Shiraishi, Y. et al. Precise characterization of somatic complex structural variations from tumor/control paired long-read sequencing data with nanomonsv. Nucleic Acids Research, gkad526 (2023).

26. Ewing, A.D. et al. Combining tumor genome simulation with crowdsourcing to benchmark somatic single-nucleotide-variant detection. Nature methods 12, 623–630 (2015).

27. Krusche, P. et al. Best practices for benchmarking germline small-variant calls in human genomes. Nature biotechnology 37, 555–560 (2019).

28. Shiraishi, Y. et al. Precise characterization of somatic complex structural variations from paired long-read sequencing data with nanomonsv. BioRxiv, 2020.2007. 2022.214262 (2020).

29. Nanopore EPI2ME Labs, https://github.com/epi2me-labs/wf-somatic-variation. (2023).

30. Sahraeian, S.M.E. et al. Achieving robust somatic mutation detection with deep learning models derived from reference data sets of a cancer sample. Genome Biology 23, 12 (2022).

31. Tarabichi, M. et al. A practical guide to cancer subclonal reconstruction from DNA sequencing. Nature methods 18, 144–155 (2021).

32. Lin, J.-H., Chen, L.-C., Yu, S.-C. & Huang, Y.-T. LongPhase: an ultra-fast chromosome-scale phasing algorithm for small and large variants. Bioinformatics 38, 1816–1822 (2022).

33. Patterson, M. et al. WhatsHap: weighted haplotype assembly for future-generation sequencing reads. Journal of Computational Biology 22, 498–509 (2015).

