## Supplementary Materials for "ClairS: a deep-learning method for long-read somatic small variant calling"

#### Supplementary Notes

Octopus (v0.7.4) ... 15
NeuSomatic (v0.2.1) ... 15
VarNet (v1.1.0) ... 16
Benchmarking ... 17
som.py (v 0.3.12) ... 17
Calculate Precision, Recall, F1-Score with different cut-off ... 17
Data availability ... 18
SEQC truth variants ... 18
GIAB truth variants ... 18
Reference genomes ... 18
Oxford Nanopore (ONT) Sequencing Data ... 19
Illumina Sequencing Data ... 19

### Supplementary Tables

#### Supplementary Table 1. Summary of datasets used for model training and testing.

**NS**: NovaSeq at Illumina; **NC**: HiSeq at National Cancer Institute; **IL**: HiSeq at Illumina; **EA**: HiSeq at European Infrastructure for Translational Medicine (EATRIS); **FD**: HiSeq at Fudan University; **NV**: HiSeq at Novartis.

| Platform | Sample | Reference | Aligner | Coverage | Source | Chemistry /Instruments | Used in training | Used for testing |
|---|---|---|---|---|---|---|---|---|
| ONT | HG001 | GRCh38_no_alt | Minimap2 | 48.44 | HKU | R10.4 | ✓ | |
| | HG002 | GRCh38_no_alt | Minimap2 | 76.29 | ONT EPI2ME Labs | R10.4 | ✓ | |
| | HCC1395 | GRCh38_no_alt | Minimap2 | 75.97 | HKU | R10.4/R10.4.1 | | ✓ |
| | HCC1395BL | GRCh38_no_alt | Minimap2 | 45.55 | HKU | R10.4/R10.4.1 | | ✓ |
| Illumina | HG003 | GRCh38_no_alt | BWA-MEM | 47.38 | Google Health Center | NovaSeq 6000 | ✓ | |
| | HG004 | GRCh38_no_alt | BWA-MEM | 46.36 | Google Health Center | NovaSeq 6000 | ✓ | |
| | HG003 | GRCh38_no_alt | BWA-MEM | 43.77 | Google Health Center | Hiseq X | ✓ | |
| | HG004 | GRCh38_no_alt | BWA-MEM | 42.13 | Google Health Center | Hiseq X | ✓ | |
| | HCC1395 | GRCh38 | BWA-MEM | 51.80 | SEQC2 (NS) | NovaSeq 6000 | | ✓ |
| | HCC1395BL | GRCh38 | BWA-MEM | 42.93 | SEQC2 (NS) | NovaSeq 6000 | | ✓ |
| | HCC1395 | GRCh38 | BWA-MEM | 39.78 | SEQC2 (NC) | Hiseq 4000 | | ✓ |
| | HCC1395BL | GRCh38 | BWA-MEM | 41.22 | SEQC2 (NC) | Hiseq 4000 | | ✓ |
| | HCC1395 | GRCh38 | BWA-MEM | 64.13 | SEQC2 (IL) | Hiseq 4000 | | ✓ |
| | HCC1395BL | GRCh38 | BWA-MEM | 56.49 | SEQC2 (IL) | Hiseq 4000 | | ✓ |
| | HCC1395 | GRCh38 | BWA-MEM | 59.43 | SEQC2 (EA) | Hiseq 4000 | | ✓ |
| | HCC1395BL | GRCh38 | BWA-MEM | 56.38 | SEQC2 (EA) | Hiseq 4000 | | ✓ |
| | HCC1395 | GRCh38 | BWA-MEM | 37.93 | SEQC2 (FD) | Hiseq 4000 | | ✓ |
| | HCC1395BL | GRCh38 | BWA-MEM | 39.45 | SEQC2 (FD) | Hiseq 4000 | | ✓ |
| | HCC1395 | GRCh38 | BWA-MEM | 86.94 | SEQC2 (NV) | Hiseq 4000 | | ✓ |
| | HCC1395BL | GRCh38 | BWA-MEM | 87.54 | SEQC2 (NV) | Hiseq 4000 | | ✓ |

#### Supplementary Table 2. Details of the ONT HCC1395/BL dataset.

| Sample | Sequencer | Chemistry | Sequencing center | Cover age | Read length N50 | Median read length | Mean read length | Mean read quality | Median read quality | Accesstion number |
|---|---|---|---|---|---|---|---|---|---|---|
| HCC1395 | PromethION 48 | R10.4 | Novogene | 58 | 20,727 | 10,604 | 12,319 | 17.8 | 17.8 | SRR25005626 |
| HCC1395 | PromethION 2 solo | R10.4.1 | HKU | 17 | 26,894 | 14,109 | 16,848 | 18.5 | 18.8 | |
| HCC1395BL | PromethION 48 | R10.4 | Novogene | 27 | 27,431 | 16,161 | 17,087 | 18.1 | 18.1 | SRR25005625 |
| HCC1395BL | PromethION 2 solo | R10.4.1 | HKU | 17 | 24,346 | 14,052 | 15,617 | 17 | 17 | |

#### Supplementary Table 3. ONT HCC1395/BL SNV calling benchmarks with multiple tumor coverages, normal coverages, and variant quality cutoffs.

ClairS and Clair3 SNV calling results of different tumor coverage, normal coverage, and variant quality cutoff combinations. Precision, Recall, F1-score, and the count of TP, FP and FN are given.

| Caller | Tumor coverage | Normal coverage | Qual cutoff | Precision | Recall | F1-score | TP | FP | FN |
|---|---|---|---|---|---|---|---|---|---|
| ClairS | 75 | 30 | 15 | 92.60% | 88.52% | 90.52% | 27,830 | 2,223 | 3,609 |
| | | 25 | | 92.94% | 86.92% | 89.83% | 27,328 | 2,076 | 4,111 |
| | | 20 | | 92.21% | 83.78% | 87.79% | 26,339 | 2,226 | 5,100 |
| | 50 | 30 | | 92.06% | 88.51% | 90.25% | 27,826 | 2,399 | 3,613 |
| | | 25 | | 93.01% | 86.86% | 89.83% | 27,309 | 2,052 | 4,130 |
| | | 20 | | 91.86% | 83.77% | 87.63% | 26,336 | 2,333 | 5,103 |
| | 25 | 30 | | 94.54% | 81.50% | 87.54% | 25,624 | 1,481 | 5,815 |
| | | 25 | | 95.03% | 78.71% | 86.11% | 24,747 | 1,293 | 6,692 |
| | | 20 | | 92.77% | 74.89% | 82.87% | 23,544 | 1,836 | 7,895 |
| | 75 | 30 | 8 | 64.14% | 96.88% | 77.18% | 30,459 | 17,032 | 980 |
| | | 25 | | 63.80% | 96.60% | 76.85% | 30,371 | 17,231 | 1,068 |
| | | 20 | | 62.66% | 96.06% | 75.85% | 30,200 | 17,996 | 1,239 |
| | 50 | 30 | | 70.56% | 96.43% | 81.49% | 30,318 | 12,652 | 1,121 |
| | | 25 | | 70.21% | 96.10% | 81.14% | 30,214 | 12,821 | 1,225 |
| | | 20 | | 68.36% | 95.69% | 79.75% | 30,083 | 13,923 | 1,356 |
| | 25 | 30 | | 83.68% | 92.07% | 87.68% | 28,947 | 5,646 | 2,492 |
| | | 25 | | 82.66% | 91.55% | 86.88% | 28,783 | 6,037 | 2,656 |
| | | 20 | | 78.35% | 91.00% | 84.20% | 28,611 | 7,908 | 2,828 |
| Clair3 | 75 | 30 | Default | 35.91% | 64.57% | 46.15% | 20,299 | 36,224 | 11,140 |
| | | 25 | | 36.21% | 64.56% | 46.40% | 20,298 | 35,755 | 11,141 |
| | | 20 | | 37.83% | 64.57% | 47.71% | 20,300 | 33,360 | 11,139 |
| | 50 | 30 | | 29.30% | 68.32% | 41.01% | 21,479 | 51,821 | 9,960 |
| | | 25 | | 29.82% | 68.32% | 41.52% | 21,478 | 50,544 | 9,961 |
| | | 20 | | 31.64% | 68.32% | 43.25% | 21,480 | 46,416 | 9,959 |
| | 25 | 30 | | 18.75% | 72.11% | 29.77% | 22,671 | 98,219 | 8,768 |
| | | 25 | | 19.27% | 72.10% | 30.42% | 22,669 | 94,944 | 8,770 |
| | | 20 | | 20.56% | 72.11% | 31.99% | 22,671 | 87,617 | 8,768 |

#### Supplementary Table 4. ONT HCC1395/BL SNV calling benchmarks with multiple tumor/normal sample purities.

ClairS and Clair3 SNV calling results of multiple tumor/normal sample purities. The tumor/normal coverage is fixed at 50x/25x. Precision, Recall, F1-score, and the count of TP, FP and FN are given.

| Caller | Tumor purity | Normal purity | QUAL cutoff | Precision | Recall | F1-score | TP | FP | FN |
|---|---|---|---|---|---|---|---|---|---|
| ClairS | 1.00 | 1.00 | 15 | 93.01% | 86.86% | 89.83% | 27,309 | 2,052 | 4,130 |
| | 0.80 | | | 96.25% | 81.63% | 88.34% | 25,663 | 999 | 5,776 |
| | 0.60 | | | 97.79% | 71.08% | 82.32% | 22,348 | 506 | 9,091 |
| | 0.40 | | | 98.68% | 52.94% | 68.91% | 16,644 | 222 | 14,795 |
| | 0.20 | | | 98.99% | 22.43% | 36.57% | 7,051 | 72 | 24,388 |
| | 1.00 | 0.95 | | 92.57% | 70.35% | 79.94% | 22,116 | 1,775 | 9,323 |
| | | 0.90 | | 91.47% | 51.74% | 66.10% | 16,267 | 1,517 | 15,172 |
| | 0.80 | 0.95 | | 95.96% | 62.74% | 75.88% | 19,726 | 830 | 11,713 |
| | 1.00 | 1.00 | 8 | 70.21% | 96.10% | 81.14% | 30,214 | 12,821 | 1,225 |
| | 0.80 | | | 80.11% | 94.45% | 86.69% | 29,694 | 7,373 | 1,745 |
| | 0.60 | | | 88.14% | 90.51% | 89.31% | 28,456 | 3,829 | 2,983 |
| | 0.40 | | | 93.81% | 80.81% | 86.83% | 25,406 | 1,675 | 6,033 |
| | 0.20 | | | 97.54% | 53.06% | 68.73% | 16,681 | 420 | 14,758 |
| | 1.00 | 0.95 | | 69.74% | 85.36% | 76.76% | 26,835 | 11,646 | 4,604 |
| | | 0.90 | | 67.01% | 69.42% | 68.19% | 21,824 | 10,746 | 9,615 |
| | 0.80 | 0.95 | | 79.68% | 81.13% | 80.40% | 25,505 | 6,505 | 5,934 |
| Clair3 | 1.00 | 1.00 | Default | 30.89% | 66.67% | 42.22% | 20,961 | 46,895 | 10,478 |
| | 0.80 | | | 28.47% | 53.10% | 37.07% | 16,693 | 41,932 | 14,746 |
| | 0.60 | | | 20.98% | 30.74% | 24.94% | 9,663 | 36,386 | 21,776 |
| | 0.40 | | | 7.07% | 7.86% | 7.45% | 2,472 | 32,475 | 28,967 |
| | 0.20 | | | 0.63% | 0.60% | 0.61% | 188 | 29,640 | 31,251 |
| | 1.00 | 0.95 | | 32.82% | 66.63% | 43.97% | 20,948 | 42,886 | 10,491 |
| | | 0.90 | | 34.34% | 66.46% | 45.28% | 20,893 | 39,945 | 10,546 |
| | 0.80 | 0.95 | | 30.63% | 53.05% | 38.84% | 16,679 | 37,767 | 14,760 |

#### Supplementary Table 5. Details of randomly chosen 300 FP and 300 FN by ClairS found in 50x/25x of ONT HCC1395/BL.

The table is in the file named "supplementary_table_5.xlsx".

#### Supplementary Table 6. ONT HCC1395/BL Indel calling benchmarks with multiple tumor coverages, normal coverages, and variant quality cutoffs.

ClairS and Clair3 Indel calling results of different tumor coverage, normal coverage, and variant quality cutoff combinations.

| Caller | Tumor coverage | Normal coverage | Qual cutoff | Precision | Recall | F1-score | TP | FP | FN |
|---|---|---|---|---|---|---|---|---|---|
| ClairS | 75 | 30 | 12 | 62.80% | 72.16% | 67.16% | 959 | 568 | 370 |
| | | 25 | | 57.79% | 71.48% | 63.91% | 950 | 694 | 379 |
| | | 20 | | 51.19% | 68.02% | 58.42% | 904 | 862 | 425 |
| | 50 | 30 | | 70.95% | 69.45% | 70.19% | 923 | 378 | 406 |
| | | 25 | | 66.54% | 66.89% | 66.72% | 889 | 447 | 440 |
| | | 20 | | 60.83% | 64.03% | 62.39% | 851 | 548 | 478 |
| | 25 | 30 | | 80.49% | 51.54% | 62.84% | 685 | 166 | 644 |
| | | 25 | | 76.91% | 49.36% | 60.13% | 656 | 197 | 673 |
| | | 20 | | 69.39% | 47.25% | 56.22% | 628 | 277 | 701 |
| | 75 | 30 | 8 | 42.19% | 76.82% | 54.47% | 1,021 | 1,399 | 308 |
| | | 25 | | 37.55% | 76.60% | 50.40% | 1,018 | 1,693 | 311 |
| | | 20 | | 32.31% | 75.32% | 45.22% | 1,001 | 2,097 | 328 |
| | 50 | 30 | | 43.77% | 76.90% | 55.79% | 1,022 | 1,313 | 307 |
| | | 25 | | 38.63% | 76.15% | 51.25% | 1,012 | 1,608 | 317 |
| | | 20 | | 33.18% | 75.70% | 46.14% | 1,006 | 2,026 | 323 |
| | 25 | 30 | | 57.96% | 66.59% | 61.97% | 885 | 642 | 444 |
| | | 25 | | 53.05% | 65.46% | 58.61% | 870 | 770 | 459 |
| | | 20 | | 44.57% | 63.96% | 52.53% | 850 | 1,057 | 479 |
| Clair3 | 75 | 30 | Default | 1.13% | 70.43% | 2.23% | 936 | 81,717 | 393 |
| | | 25 | | 1.05% | 70.05% | 2.06% | 931 | 88,024 | 398 |
| | | 20 | | 0.93% | 70.13% | 1.84% | 932 | 99,306 | 397 |
| | 50 | 30 | | 1.07% | 71.41% | 2.11% | 949 | 87,787 | 380 |
| | | 25 | | 1.00% | 71.11% | 1.97% | 945 | 93,631 | 384 |
| | | 20 | | 0.90% | 71.03% | 1.78% | 944 | 104,046 | 385 |
| | 25 | 30 | | 0.79% | 69.53% | 1.57% | 924 | 115,426 | 405 |
| | | 25 | | 0.76% | 69.15% | 1.50% | 919 | 119,980 | 410 |
| | | 20 | | 0.71% | 69.15% | 1.40% | 919 | 128,637 | 410 |

#### Supplementary Table 7. ClairS ablation study.

An ablation study of ClairS. The tumor/normal coverage is fixed at 50x/25x. Prioritize-recall mode was used. Precision, Recall, F1-score, and the count of TP, FP and FN are given.

| Experiments | Precision | Recall | F1-score | TP | FP | FN |
|---|---|---|---|---|---|---|
| Default | 70.21% | 96.10% | 81.14% | 30,214 | 12,821 | 1,225 |
| Disable tumor read phasing | 66.50% | 96.27% | 78.66% | 30,267 | 15,249 | 1,172 |
| Pileup network only | 58.98% | 96.71% | 73.27% | 30,404 | 21,145 | 1,035 |
| Full-alignment network only | 67.28% | 96.05% | 79.14% | 30,198 | 14,683 | 1,241 |
| Disable haplotype consistency | 67.14% | 96.38% | 79.14% | 30,301 | 14,832 | 1,138 |

**Supplementary Table 8. Illumina HCC1395/BL SNV calling benchmarks with different somatic variant callers.**

The Precision, Recall, F1-score, and the count of TP, FP and FN of HCC1395/BL short-read datasets from six SEQC2 sources (NS: NovaSeq at Illumina, NC: HiSeq at National Cancer Institute, IL: HiSeq at Ilumina, EA: HiSeq European Infrastructure for Translational Medicine, FD: HiSeq Fudan University, NV: HiSeq Novartis) using seven somatic variant callers (Strelka2, Lancet, Mutect2, Neusomatic, Octopus, Varnet, ClairS).

| Source | Sequencer | Caller | Precision | Recall | F1 | TP | FP | FN |
|---|---|---|---|---|---|---|---|---|
| NS | NovaSeq 6000 | ClairS | 98.04% | 97.72% | 97.88% | 36,180 | 722 | 843 |
| NS | NovaSeq 6000 | Strelka2 | 95.16% | 97.18% | 96.16% | 35,978 | 1,829 | 1,045 |
| NS | NovaSeq 6000 | Mutect2 | 98.32% | 92.29% | 95.21% | 34,170 | 583 | 2,853 |
| NS | NovaSeq 6000 | Lancet | 98.47% | 85.52% | 91.54% | 31,663 | 491 | 5,360 |
| NS | NovaSeq 6000 | Neusomatic | 77.39% | 93.51% | 84.69% | 34,620 | 10,113 | 2,403 |
| NS | NovaSeq 6000 | Octopus | 78.27% | 87.13% | 82.46% | 32,259 | 8,957 | 4,764 |
| NS | NovaSeq 6000 | SomaticSniper | 58.45% | 84.50% | 69.10% | 31,286 | 22,239 | 5,737 |
| NS | NovaSeq 6000 | Varnet | 98.94% | 85.92% | 91.97% | 31,812 | 342 | 5,211 |
| NC | HiSeq 4000 | ClairS | 98.82% | 96.84% | 97.82% | 35,109 | 418 | 1,145 |
| NC | HiSeq 4000 | Strelka2 | 95.81% | 96.55% | 96.18% | 35,002 | 1,530 | 1,252 |
| NC | HiSeq 4000 | Mutect2 | 98.58% | 93.08% | 95.75% | 33,744 | 485 | 2,510 |
| NC | HiSeq 4000 | Lancet | 99.45% | 89.48% | 94.20% | 32,439 | 180 | 3,815 |
| NC | HiSeq 4000 | Neusomatic | 90.34% | 95.21% | 92.72% | 34,519 | 3,689 | 1,735 |
| NC | HiSeq 4000 | Octopus | 90.85% | 83.62% | 87.09% | 30,315 | 3,052 | 5,939 |
| NC | HiSeq 4000 | SomaticSniper | 58.00% | 86.98% | 69.59% | 31,532 | 22,837 | 4,722 |
| NC | HiSeq 4000 | Varnet | 98.83% | 85.78% | 91.84% | 31,097 | 368 | 5,157 |
| IL | HiSeq 4000 | ClairS | 98.93% | 96.04% | 97.46% | 35,843 | 388 | 1,478 |
| IL | HiSeq 4000 | Strelka2 | 96.74% | 97.33% | 97.03% | 36,323 | 1,225 | 998 |
| IL | HiSeq 4000 | Mutect2 | 98.71% | 92.18% | 95.33% | 34,403 | 449 | 2,918 |
| IL | HiSeq 4000 | Lancet | 99.39% | 91.54% | 95.31% | 34,165 | 210 | 3,156 |
| IL | HiSeq 4000 | Neusomatic | 92.14% | 93.72% | 92.93% | 34,979 | 2,982 | 2,342 |
| IL | HiSeq 4000 | Octopus | 86.99% | 84.57% | 85.76% | 31,561 | 4,719 | 5,760 |
| IL | HiSeq 4000 | SomaticSniper | 55.01% | 86.63% | 67.29% | 32,332 | 26,444 | 4,989 |
| IL | HiSeq 4000 | Varnet | 98.69% | 89.26% | 93.74% | 33,313 | 443 | 4,008 |
| EA | HiSeq 4000 | ClairS | 98.86% | 96.23% | 97.53% | 35,872 | 412 | 1,404 |
| EA | HiSeq 4000 | Strelka2 | 96.54% | 98.18% | 97.35% | 36,597 | 1,313 | 679 |
| EA | HiSeq 4000 | Mutect2 | 98.48% | 94.09% | 96.24% | 35,072 | 540 | 2,204 |
| EA | HiSeq 4000 | Lancet | 99.30% | 90.85% | 94.89% | 33,865 | 240 | 3,411 |
| EA | HiSeq 4000 | Neusomatic | 86.86% | 95.40% | 90.93% | 35,560 | 5,378 | 1,716 |
| EA | HiSeq 4000 | Octopus | 89.19% | 84.76% | 86.92% | 31,597 | 3,831 | 5,679 |
| EA | HiSeq 4000 | SomaticSniper | 56.53% | 86.56% | 68.40% | 32,265 | 24,807 | 5,011 |
| EA | HiSeq 4000 | Varnet | 98.96% | 91.14% | 94.89% | 33,972 | 358 | 3,304 |
| FD | HiSeq 4000 | ClairS | 97.57% | 96.50% | 97.03% | 35,005 | 872 | 1,270 |
| FD | HiSeq 4000 | Strelka2 | 95.65% | 97.00% | 96.32% | 35,186 | 1,600 | 1,089 |
| FD | HiSeq 4000 | Mutect2 | 98.68% | 94.34% | 96.46% | 34,223 | 459 | 2,052 |
| FD | HiSeq 4000 | Lancet | 99.43% | 90.11% | 94.54% | 32,686 | 187 | 3,589 |
| FD | HiSeq 4000 | Neusomatic | 77.62% | 95.71% | 85.72% | 34,718 | 10,012 | 1,557 |
| FD | HiSeq 4000 | Octopus | 92.94% | 83.60% | 88.02% | 30,327 | 2,304 | 5,948 |
| FD | HiSeq 4000 | SomaticSniper | 59.97% | 87.08% | 71.03% | 31,589 | 21,085 | 4,686 |
| FD | HiSeq 4000 | Varnet | 98.92% | 87.70% | 92.97% | 31,813 | 348 | 4,462 |
| NV | HiSeq 4000 | ClairS | 98.85% | 94.10% | 96.41% | 35,432 | 414 | 2,221 |
| NV | HiSeq 4000 | Strelka2 | 95.83% | 95.11% | 95.47% | 35,813 | 1,559 | 1,840 |
| NV | HiSeq 4000 | Mutect2 | 99.02% | 90.09% | 94.35% | 33,923 | 336 | 3,730 |
| NV | HiSeq 4000 | Lancet | 99.43% | 89.31% | 94.10% | 33,628 | 192 | 4,025 |
| NV | HiSeq 4000 | Neusomatic | 92.46% | 92.86% | 92.66% | 34,965 | 2,852 | 2,688 |
| NV | HiSeq 4000 | Octopus | 84.07% | 84.95% | 84.51% | 31,987 | 6,061 | 5,666 |
| NV | HiSeq 4000 | SomaticSniper | 52.19% | 85.68% | 64.86% | 32,260 | 29,557 | 5,393 |
| NV | HiSeq 4000 | Varnet | 99.09% | 90.11% | 94.38% | 33,929 | 313 | 3,724 |

### Supplementary Methods

#### Description of pileup input features

Pileup input comprises 34 features at each genome position. The features provide information on the counts of nucleotides, insertions, deletions, low mapping quality, and low-quality bases in both the forward and reverse strands. Here's a breakdown of each channel:

1-4: A+/C+/G+/T+: These features record the counts of A/C/G/T nucleotides present in the forward strand.

5: $I_S$+: This channel provides the count of insertions that have the same starting positions as the candidate site in the forward strand.

6: $I_{1S}$+: Similar to $I_S$+ but only counting the insertion with the highest read support.

7: $D_S$+: This channel provides the count of deletions that have  the same starting positions as the candidate site in the forward strand.

8: $D_{1S}$+: Similar to $D_S$+ but only counting the deletion with the highest read support.

9: $D_R$+: The count of all deletions in the forward strand that crossed the position except for the first base of each deletion.

10-13: A-/C-/G-/T-: These features record the counts of A/C/G/T nucleotides present in the reverse strand.

14: $I_S$ This channel provides the count of insertions that have the same starting positions as the candidate site in the reverse strand.

15: $I_{1S}$-: Similar to $I_S$- but only counting the insertion with the highest read support.

16: $D_S$-: This channel provides the count of deletions that have  the same starting positions as the candidate site in the reverse strand.

17: $D_{1S}$-: Similar to $D_S$- but only counting the deletion with the highest read support.

18: $D_R$-: The count of all deletions in the reverse strand that crossed the position except for the first base of each deletion.

19-22: $A_{LMQ}$+/$C_{LMQ}$+/$G_{LMQ}$+/$T_{LMQ}$+: Theses features record the counts of A/C/G/T nucleotides with low mapping quality (MQ < 20) in the forward strand.

23-26: $A_{LBQ}$+/$C_{LBQ}$+/$G_{LBQ}$+/$T_{LBQ}$+: These features record the counts of A/C/G/T nucleotides with low base quality (BQ < 30) in the forward strand.

27-30: $A_{LMQ}$-/$C_{LMQ}$-/$G_{LMQ}$-/$T_{LMQ}$-: These features record the counts of A/C/G/T nucleotides with low mapping quality (MQ < 20) in the reverse strand.

31-34: $A_{LBQ}$-/$C_{LBQ}$-/$G_{LBQ}$-/$T_{LBQ}$-: These features record the counts of A/C/G/T nucleotides with low base quality (BQ < 30) in the reverse strand.

#### Description of full-alignment input channels

Full-alignment input comprises seven channels. Each channel is a two-dimensional representation (reads as rows, and genome positions as columns) of a type of information extracted from the read alignments. Here's a breakdown of each channel:

1: Reference base: This channel assigned an integer {A:100, G:75, T:50, C:25} to each position based on the reference base. Deleted bases are assigned 0.

2: Alternative base: Positions where the alignment mismatches the reference base are assigned an integer value {A:100, G:75, T:50, C:25, insertion starting position: -50, deletion: -100}. If a position matches the reference base, 0 is assigned.

3: Strand information: All positions of a read are assigned an integer value {+: 100, -: -100} based on the strand the read aligns to. For positions in a deletion, 0 is assigned.

4: Base quality: Each aligned base is assigned an integer value ranging from 0 to 100, scaled up from the Phred base quality score (0 to 60). If the score exceeds 60, it is capped at 60. Deleted bases are assigned 0. Decimal places are truncated.

5: Mapping quality: The whole read is assigned an integer value ranging from 0 to 100, scaled up from the Phred mapping quality (0 to 60) of the aligned read. The Phred mapping quality (MQ) is rounded as follows: {MQ<20:20, 20≤MQ<40: 40, MQ≥40: 60}. Deleted bases are assigned 0. Decimal places are truncated.

6: Tumor/Normal/Phasing information: For reads from the tumor sample, an integer value {HP1: 30, unphased: 60, HP2: 90} is assigned to each position of the read. For reads from the normal sample, an integer value -60 is assigned. Deleted bases are assigned 0. If the reads are phased, the tumor reads in all seven channels are sorted in the order "unphased, HP1, HP2."

7: Insertion base: Inserted bases are exhibited after each insertion's starting position with integers {A:100, G:75, T:50, C:25}.

### Command lines used

#### Read alignment

##### Minimap2 (v 2.17-r941)

```
# Align ONT reads using minimap2 to GRCh38_no_alt by default
minimap2 -t ${THREADS} -aL -z 600,200 -x map-ont ref.fa input.fastq.gz | samtools view -bh
-o output.unsorted.bam -
samtools sort -@${THREADS} -o output.sorted.bam output.unsorted.bam && samtools index -@
${THREADS} output.sorted.bam
```

#### BAM subsampling

##### Samtools(v 1.15.1)

```
samtools view -@ ${THREADS} -s ${RATIO}.${RATIO} -b -o subsampled.bam ${BAM}
samtools index -@ ${THREADS} subsampled.bam
```

#### Coverage calculation

##### Mosdepth(v 0.3.1)

```
mosdepth -t ${THREADS} -n -x --quantize 0:15:150: output ${BAM}
```

#### Alignment statistical summary

##### NanoPlot(v 1.40.2)

```
NanoPlot -t ${THREADS} --bam ${BAM} --N50 -o bamplots
```

#### Generating BAMs with different tumor/normal purities

```
# Use ${TUMOR_PURITY} to add tumor purity
# Use ${NORMAL_PURITY} to add normal purity
pypy3 clairs.py gen_contaminated_bam \
    --tumor_bam_fn ${TUMOR_BAM_FILE_PATH} \
    --normal_bam_fn ${NORMAL_BAM_FILE_PATH} \
    --tumor_purity ${TUMOR_PURITY} \
    --normal_purity ${NORMAL_PURITY} \
    --output_dir ${OUTPUT_DIR} \
    --tumor_bam_coverage ${TUMOR_BAM_COVERAGE} \
    --normal_bam_coverage ${NORMAL_BAM_COVERAGE} \
    --mosdepth ${MOSDEPTH_PATH}
```

#### Running ClairS (v0.0.1)

```
run_clairs \
  --tumor_bam_fn ${TUMOR_BAM_FILE_PATH} \
```

```
  --normal_bam_fn ${NORMAL_BAM_FILE_PATH} \
  --ref_fn ${REF} \
  --threads ${THREADS} \
  --platform ${PLATFORM} \
  --output ${OUTPUT_DIR}
```

#### Running Clair3 (v0.1-r12)

```
# The Clair3 R10.4 Q20+ model was downloaded from https://github.com/nanoporetech/rerio
# maintained by ONT developer
# Then run the following command in both tumor and normal BAMs
docker run -it \
  -v ${INPUT_DIR}:${INPUT_DIR} \
  -v ${OUTPUT_DIR}:${OUTPUT_DIR} \
  hkubal/clair3:v0.1-r12 \
  /opt/bin/run_clair3.sh \
  --bam_fn=${INPUT_DIR}/input.bam \
  --ref_fn=${INPUT_DIR}/ref.fa \
  --threads=${THREADS} \
  --platform="${PLATFORM}" \
  --model_path="${INPUT_DIR}/r104_e81_sup_g5015" \
  --output=${OUTPUT_DIR}

# Acquire somatic variants in tumor and normal output
pypy3 clairs.py clair3_somatic_calling \
  --tumor_vcf_fn clair3_tumor.vcf.gz \
  --normal_vcf_fn clair3_normal.vcf.gz \
  --output_vcf_fn clair3_output.vcf.gz
```

#### Running other somatic variant callers for Illumina data

##### Strelka2 (v2.9.10)

```
configureStrelkaSomaticWorkflow.py \
    --normalBam ${NORMAL_BAM_FILE_PATH} \
    --tumorBam ${TUMOR_BAM_FILE_PATH} \
    --referenceFasta ${REF} \
    --retainTempFiles \
    --runDir ${OUTPUT_DIR}/demo_somatic \
    --callRegions ${BED_FILE_PATH}

# execution with ${threads} parallel jobs
${OUTPUT_FOLDER}/demo_somatic/runWorkflow.py -m local -j ${threads}
```

##### Mutect2 (v4.2.6.1)

```
gatk --java-options -Xmx12G Mutect2 \
```

```
    --reference ${REF} \
    --intervals ${BED_FILE_PATH} \
    --input ${NORMAL_BAM_FILE_PATH} \
    --input ${TUMOR_BAM_FILE_PATH} \
    --normal ${NORMAL_SAMPLE} \
    --output ${OUTPUT_DIR}/unfiltered.mutect2.vcf

gatk --java-options -Xmx12G FilterMutectCalls \
    --variant ${OUTPUT_DIR}/unfiltered.mutect2.vcf \
    --reference ${REF} \
    --output ${OUTPUT_DIR}/mutect2.vcf)
```

#### Lancet (v4.2.6.1)

```
NUMBER_OF_AUTOSOMES=22
for chrom in `seq 1 $NUMBER_OF_AUTOSOMES`
do
    mkdir -p ${OUTPUT_FOLDER}/${chrom}
    lancet \
           --tumor ${TUMOR_BAM_FILE_PATH} \
           --normal ${NORMAL_BAM_FILE_PATH} \
           --ref ${REF} \
           -B ${INPUT_DIR}/chr${chrom}.bed \
           --num-threads ${THREADS} > ${OUTPUT_DIR}/${chrom}/output.vcf
done
# concat all VCF records
vcf-concat ${OUTPUT_DIR}/*/output.vcf | vcf-sort > ${OUTPUT_DIR}/output.vcf
```

#### Octopus (v0.7.4)

```
octopus \
    -R ${REF} \
    -I ${NORMAL_BAM_FILE_PATH} ${TUMOR_BAM_FILE_PATH} \
    -N ${NORMAL_SAMPLE} \
    --somatics-only \
    --threads ${THREADS} \
    -o ${OUTPUT_FOLDER}/output.vcf
```

#### NeuSomatic (v0.2.1)

```
python neusomatic/python/preprocess.py \
    --mode call \
    --reference ${REF} \
    --region_bed ${BED_FILE_PATH} \
    --tumor_bam ${TUMOR_BAM_FILE_PATH} \
    --normal_bam ${NORMAL_BAM_FILE_PATH} \
```

```
    --work ${OUTPUT_DIR}/work_standalone \
    --min_mapq 10 \
    --num_threads ${THREADS} \
    --scan_alignments_binary neusomatic/bin/scan_alignments

python neusomatic/python/call.py \
       --candidates_tsv ${OUTPUT_DIR}/work_standalone/dataset/*/candidates*.tsv \
       --reference ${REF} \
       --out ${OUTPUT_DIR} \
       --checkpoint ${MODEL_PATH} \
       --num_threads ${THREADS} \
       --batch_size 100

python ${NS_PATH}/neusomatic/python/postprocess.py \
       --reference ${REF} \
       --tumor_bam ${TUMOR_BAM_FILE_PATH} \
       --pred_vcf ${OUTPUT_DIR}/pred.vcf \
       --candidates_vcf ${OUTPUT_DIR}/work_standalone/work_tumor/filtered_candidates.vcf \
       --output_vcf ${OUTPUT_DIR}/NeuSomatic_standalone.vcf \
       --work ${OUTPUT_DIR}
```

#### VarNet (v1.1.0)

```
python VarNet/filter.py \
       --sample_name ${sample_name} \
       --normal_bam ${NORMAL_BAM_FILE_PATH} \
       --tumor_bam ${TUMOR_BAM_FILE_PATH} \
       --processes ${THREADS} \
       --output_dir ${OUTPUT_DIR} \
       --reference ${REF} \
       --region_bed ${BED_FILE_PATH} \
       -snv

python VarNet/predict.py \
       --sample_name ${sample_name} \
       --normal_bam ${NORMAL_BAM_FILE_PATH} \
       --tumor_bam ${TUMOR_BAM_FILE_PATH} \
       --processes ${THREADS} \
       --output_dir ${OUTPUT_DIR} \
       --reference ${REF} \
       -snv
```

#### Benchmarking

##### som.py (v 0.3.12)

```
som.py ${SEQC2_BASELINE_VCF} output.vcf.gz \
       -T ${SEQC2_CONFIDENT_BED} \
       -f ${SEQC2_CONFIDENT_BED} \
       -r ${REF} \
       -o benchmark_result
```

##### Calculate Precision, Recall, F1-Score with different cut-off

```
pypy3 clairs.py compare_vcf \
       --truth_vcf_fn ${SEQC2_BASELINE_VCF} \
       --input_vcf_fn output.vcf.gz \
       --bed_fn ${SEQC2_CONFIDENT_BED \
       --output_dir benchmark_result \
       --input_filter_tag 'PASS' \
       --min_qual ${MIN_QUAL} \
       --min_af ${MIN_AF} \
       --output_best_f1_score \
       --tumor_bam_fn ${TUMOR_BAM_FILE_PATH} \
       --normal_bam_fn ${NORMAL_BAM_FILE_PATH}
```

### Data availability

#### SEQC truth variants

**HCC1395-HCC1395BL pair, GRCh38, v1.2**
https://ftp-trace.ncbi.nlm.nih.gov/ReferenceSamples/seqc/Somatic_Mutation_WG/release/v1.2

#### GIAB truth variants

**HG001 (NA12878), GRCh38, v4.2.1**
https://ftp-trace.ncbi.nlm.nih.gov/giab/ftp/release/NA12878_HG001/NISTv4.2.1/GRCh38/

**HG002 (NA24385), GRCh38, v 4.2.1**
https://ftp-trace.ncbi.nlm.nih.gov/giab/ftp/release/AshkenazimTrio/HG002_NA24385_son/NISTv4.2.1/GRCh38/

**HG003 (NA24149), GRCh38, v 4.2.1**
https://ftp-trace.ncbi.nlm.nih.gov/giab/ftp/release/AshkenazimTrio/HG003_NA24149_father/NISTv4.2.1/GRCh38/

**HG004 (NA24143), GRCh38, v 4.2.1**
https://ftp-trace.ncbi.nlm.nih.gov/giab/ftp/release/AshkenazimTrio/HG004_NA24143_mother/NISTv4.2.1/GRCh38/

#### Reference genomes

**GRCh38 d1.vd1**
https://ftp-trace.ncbi.nlm.nih.gov/ReferenceSamples/seqc/Somatic_Mutation_WG/technical/reference_genome/GRCh38/GRCh38.d1.vd1.fa

**GRCh38_no_alt**
https://ftp.ncbi.nlm.nih.gov/genomes/all/GCA/000/001/405/GCA_000001405.15_GRCh38/seqs_for_alignment_pipelines.ucsc_ids/GCA_000001405.15_GRCh38_no_alt_analysis_set.fna.gz

**GRCh38 Stratification regions (v 2.0)**
https://ftp-trace.ncbi.nlm.nih.gov/giab/ftp/release/genome-stratifications/v2.0/GRCh38

#### Oxford Nanopore (ONT) Sequencing Data

**HCC1395 R10.4.1 Q20+, GRCh38_no_alt, 75.91-fold**
https://www.ncbi.nlm.nih.gov/sra/?term=SRR25005626

**HCC1395BL R10.4.1 Q20+, GRCh38_no_alt, 45.55-fold**
https://www.ncbi.nlm.nih.gov/sra/?term=SRR25005625

**HG001 R10.4.1 Q20+ (NA12878), GRCh38_no_alt, 48.44-fold**
http://www.bio8.cs.hku.hk/clairs/data/ONT_HG001_R10.4_GRch38.bam

**HG002 R10.4 Q20+ (NA24385) , GRCh38_no_alt, 76.29-fold**
https://labs.epi2me.io/gm24385_q20_2021.10

#### Illumina Sequencing Data

**HCC1395 NovaSeq 6000 (SEQC2 NS), GRCh38, 51.80-fold**
https://ftp-trace.ncbi.nlm.nih.gov/ReferenceSamples/seqc/Somatic_Mutation_WG/data/WGS/WGS_NS_T_1.bwa.dedup.bam

**HCC1395BL NovaSeq 6000 (SEQC2 NS), GRCh38, 42.93-fold**
https://ftp-trace.ncbi.nlm.nih.gov/ReferenceSamples/seqc/Somatic_Mutation_WG/data/WGS/WGS_NS_N_1.bwa.dedup.bam

**HCC1395 Hiseq 4000 (SEQC2 EA), GRCh38, 59.43-fold**
https://ftp-trace.ncbi.nlm.nih.gov/ReferenceSamples/seqc/Somatic_Mutation_WG/data/WGS/WGS_EA_T_1.bwa.dedup.bam

**HCC1395BL Hiseq 4000 (SEQC2 EA), GRCh38, 56.38-fold**
https://ftp-trace.ncbi.nlm.nih.gov/ReferenceSamples/seqc/Somatic_Mutation_WG/data/WGS/WGS_EA_N_1.bwa.dedup.bam

**HCC1395 Hiseq 4000 (SEQC2 FD), GRCh38, 37.93-fold**
https://ftp-trace.ncbi.nlm.nih.gov/ReferenceSamples/seqc/Somatic_Mutation_WG/data/WGS/WGS_FD_T_1.bwa.dedup.bam

**HCC1395BL Hiseq 4000 (SEQC2 FD), GRCh38, 39.45-fold**
https://ftp-trace.ncbi.nlm.nih.gov/ReferenceSamples/seqc/Somatic_Mutation_WG/data/WGS/WGS_FD_N_1.bwa.dedup.bam

**HCC1395 Hiseq 4000 (SEQC2 IL), GRCh38, 64.13-fold**
https://ftp-trace.ncbi.nlm.nih.gov/ReferenceSamples/seqc/Somatic_Mutation_WG/data/WGS/WGS_IL_T_1.bwa.dedup.bam

**HCC1395BL Hiseq 4000 (SEQC2 IL), GRCh38, 56.49-fold**
https://ftp-trace.ncbi.nlm.nih.gov/ReferenceSamples/seqc/Somatic_Mutation_WG/data/WGS/WGS_IL_N_1.bwa.dedup.bam

**HCC1395 Hiseq 4000 (SEQC2 LL), GRCh38, 38.84-fold**
https://ftp-trace.ncbi.nlm.nih.gov/ReferenceSamples/seqc/Somatic_Mutation_WG/data/WGS/WGS_LL_T_1.bwa.dedup.bam

**HCC1395BL Hiseq 4000 (SEQC2 LL), GRCh38, 39.16-fold**
https://ftp-trace.ncbi.nlm.nih.gov/ReferenceSamples/seqc/Somatic_Mutation_WG/data/WGS/WGS_LL_N_1.bwa.dedup.bam

**HCC1395 Hiseq 4000 (SEQC2 NC), GRCh38, 39.78-fold**
https://ftp-trace.ncbi.nlm.nih.gov/ReferenceSamples/seqc/Somatic_Mutation_WG/data/WGS/WGS_NC_T_1.bwa.dedup.bam

**HCC1395BL Hiseq 4000 (SEQC2 NC), GRCh38, 41.22-fold**
https://ftp-trace.ncbi.nlm.nih.gov/ReferenceSamples/seqc/Somatic_Mutation_WG/data/WGS/WGS_NC_N_1.bwa.dedup.bam

**HCC1395 Hiseq 4000 (SEQC2 NV), GRCh38, 86.94-fold**
https://ftp-trace.ncbi.nlm.nih.gov/ReferenceSamples/seqc/Somatic_Mutation_WG/data/WGS/WGS_NV_T_1.bwa.dedup.bam

**HCC1395BL Hiseq 4000 (SEQC2 NV), GRCh38, 87.54-fold**
https://ftp-trace.ncbi.nlm.nih.gov/ReferenceSamples/seqc/Somatic_Mutation_WG/data/WGS/WGS_NV_N_1.bwa.dedup.bam

**HG003 NovaSeq 6000 (NA24149), GRCh38_no_alt, 47.38-fold**
https://storage.googleapis.com/brain-genomics-public/research/sequencing/grch38/bam/novaseq/wgs_pcr_free/50x/HG003.novaseq.pcr-free.50x.dedup.grch38.bam

**HG003 HiSeqX (NA24149), GRCh38_no_alt 43.77-fold**

https://storage.googleapis.com/brain-genomics-public/research/sequencing/grch38/bam/hiseqx/wgs_pcr_free/40x/HG003.hiseqx.pcr-free.40x.dedup.grch38.bam

**HG004 NovaSeq 6000 (NA24385), GRCh38_no_alt, 46.36-fold**
https://storage.googleapis.com/brain-genomics-public/research/sequencing/grch38/bam/novaseq/wgs_pcr_free/50x/HG004.novaseq.pcr-free.50x.dedup.grch38.bam

**HG004 HiSeqX (NA24385), GRCh38_no_alt, 42.13-fold**
https://storage.googleapis.com/brain-genomics-public/research/sequencing/grch38/bam/hiseqx/wgs_pcr_free/40x/HG004.hiseqx.pcr-free.40x.dedup.grch38.bam
